# TEAD switches interacting partners along neural progenitor lineage progression to execute distinct functions

**DOI:** 10.1101/2024.12.19.629472

**Authors:** Charles H. Perry, Alfonso Lavado, Venkata Thulabandu, Cody Ramirez, Joshua Paré, Rajiv Dixit, Akhilesh Mishra, Jiyuan Yang, Jiyang Yu, Xinwei Cao

## Abstract

The TEAD family of transcription factors are best known as the DNA-binding factor in the Hippo pathway, where they act by interacting with transcriptional coactivators YAP and TAZ (YAP/TAZ). Despite the importance of the Hippo pathway, the in vivo functions of TEAD in mammals have not been well established. By comparing mouse mutants lacking TEAD1 and TEAD2 (TEAD1/2) to those lacking YAP/TAZ, we found that TEAD1/2 have both YAP/TAZ-dependent and -independent functions during ventral telencephalon development. TEAD1/2 loss and YAP/TAZ loss similarly disrupt neuroepithelial apical junctions. However, the impacts of their losses on progenitor lineage progression are essentially opposite: Whereas YAP/TAZ loss depletes early progenitors and increases later progenitors—consistent with their established function in promoting progenitor self-renewal and proliferation, TEAD1/2 loss expands early progenitors and reduces late progenitors, indicating that TEAD1/2 promote lineage progression. We further show that TEAD1/2 promote neural progenitor lineage progression by, at least in part, inhibiting Notch signaling and by cooperating with Insulinoma-associated 1 (INSM1). Orthologs of TEAD and INSM1 have been shown to cooperatively regulate neuronal cell fate decisions in worms and flies. Our study reveals a remarkable evolutionary conservation of the function of this transcription factor complex during metazoan neural development.

## Introduction

In the developing central nervous system (CNS), the primary stem/progenitor cells, neuroepithelial cells (NECs) and the cells they transform into upon the onset of neurogenesis— radial glia (RG), form the ventricular zone (VZ) lining the central lumen of the neural tube (Kriegstein and Alvarez-Buylla 2009; Taverna et al. 2014). These cells divide—depending on their division mode—to expand or replenish themselves, produce neurons directly (direct neurogenesis), or give rise to another type of progenitor cells known as intermediate progenitors (IPs). IPs, residing in the subventricular zone (SVZ) adjacent to the VZ, divide to expand themselves and produce neurons (indirect neurogenesis), increasing the cell output of primary stem/progenitor cells. NECs/RG and IPs exhibit key cell biological differences. NECs/RG undergo mitosis at or near the ventricular surface (VS); they are also called apical progenitors (APs). IPs undergo mitosis in the SVZ; they are also called basal progenitors (BPs). Moreover, it is generally thought that RG are multipotent (being able to produce different types of neurons or both neurons and glia), whereas IPs are unipotent (being able to produce only one type of neurons), and that RGs divide more rounds than IPs.

Our knowledge about RG and IPs is mostly obtained by studying progenitor cells in the developing mouse cortex. Neural progenitors in the ventral telencephalon (subpallium), which give rise to extremely diverse types of cells including not only neurons and glia that form the basal ganglia, parts of the septum and the amygdala, the preoptic area, but also all cortical interneurons and a subset of cortical oligodendrocyte lineage cells (OLCs) (Anderson et al. 1997; Kessaris et al. 2006; Moreno et al. 2009; Turrero Garcia and Harwell 2017), are likely to possess unique features that underlie their distinct cellular output. For example, whereas the vast majority of progenitor cells reside in the VZ in most regions of the developing mouse CNS, in the ganglionic eminences (GEs, subregions of the subpallium) the SVZ largely surpasses the VZ in size and is the main site of cell proliferation throughout most of the neurogenic period (Smart 1976; Fentress et al. 1981; Pilz et al. 2013). Furthermore, studies have found novel subtypes of progenitor cells (e.g., subapical progenitors (SAPs), which are common in the mouse GEs and in the developing gyrated cerebral cortices of ferrets and sheep but rare in the developing mouse cortex) and distinct progenitor behaviors (e.g., more proliferative divisions by several subtypes of progenitor cells and a progressively faster cell cycle) in the subpallium (Pilz et al. 2013). It remains largely unknown what mechanisms uniquely govern the characteristics and lineage progression of subpallial neural progenitors.

The Hippo pathway plays important roles in the development, tumorigenesis, and regeneration of many tissues across species (Zheng and Pan 2019). It controls the activity of a transcription factor complex composed of the transcriptional coactivator YAP (Yorki in flies, YAP and TAZ (YAP/TAZ) in mammals) and the DNA-binding factor TEAD (Scalloped in flies, TEAD1–4 in mammals). In many tissues, YAP is expressed in the stem/progenitor cells and promotes their proliferation and self-renewal (Driskill and Pan 2023; Zhong et al. 2024). In contrast to the extensive studies on the functions of YAP/TAZ, studies on the in vivo function of TEAD in mammals are scarce (Landin-Malt et al. 2016; Lin et al. 2017), probably because it is assumed that TEAD simply acts as the DNA-binding partner of YAP/TAZ. Some observations, however, suggest that TEAD acts with other cofactors in some contexts. TEAD is evolutionarily much more ancient than YAP; TEAD is present in amoebozoans whereas YAP appears to be restricted to holozoans (Sebé-Pedrós et al. 2012; Phillips et al. 2024). In *Drosophila*, Scalloped, identified in 1929 as a result of a wing phenotype (Grunberg 1929), regulates wing identity independent of Yorki/YAP (Halder et al. 1998; Simmonds et al. 1998). In *C. elegans*, the TEAD ortholog EGL-44 regulates neuron cell fate and neuroblast lineage progression in an apparently YAP-independent manner (Wu et al. 2001; Feng et al. 2013; Iwasa et al. 2013). We have reported that, in the developing chick spinal cord, YAP is only expressed in the VZ but TEAD is expressed not only in the VZ but also in nearby cells expressing the proneural transcription factor Neurogenin 2 (Cao et al. 2008), suggesting that TEAD may have YAP-independent function in these Neurogenin 2-expressing cells.

In this study, we show that TEAD has both YAP/TAZ-dependent and -independent functions during mammalian ventral telencephalon development. In APs/RG, TEAD acts with YAP/TAZ to maintain the structural integrity of neuroepithelial apical junctions. In IPs, TEAD acts with Insulinoma-associated 1 (INSM1) to promotes lineage progression at least in part by inhibiting Notch signaling.

## Results

### TEAD1 is expressed in broader regions than YAP/TAZ are in the developing telencephalon

Our earlier observation in the chick neural tube prompted us to examine YAP/TAZ and TEAD expression patterns in the developing mammalian telencephalon. We compared YAP/TAZ and TEAD1 immunostaining patterns to those of SOX2, which marks RG and some IPs throughout the developing CNS; TBR2, which marks cortical IPs (Englund et al. 2005); and ASCL1, which marks subsets of subpallial neural progenitors (Yun et al. 2002; Petryniak et al. 2007; Imayoshi et al. 2013a) (Fig. 1A). In the cortex, as we have reported previously (Lavado et al. 2021), YAP/TAZ were specifically expressed in SOX2^+^TBR2^−^ RG. TEAD1 immunosignals, however, were detected in the VZ, the TBR2-dense SVZ, and weakly above the SVZ, suggesting that TEAD1 is expressed in both RG and IPs, and probably also in new-born neurons. In the medial ganglionic eminence (MGE), a subregion of the subpallium, YAP/TAZ were expressed in the cells near the VS, presumably RG, whereas TEAD1 was detected in both the VZ and the adjacent SVZ. YAP/TAZ and TEAD1 showed similar expression patterns in the lateral and caudal GEs (data not shown).

**Figure 1.**
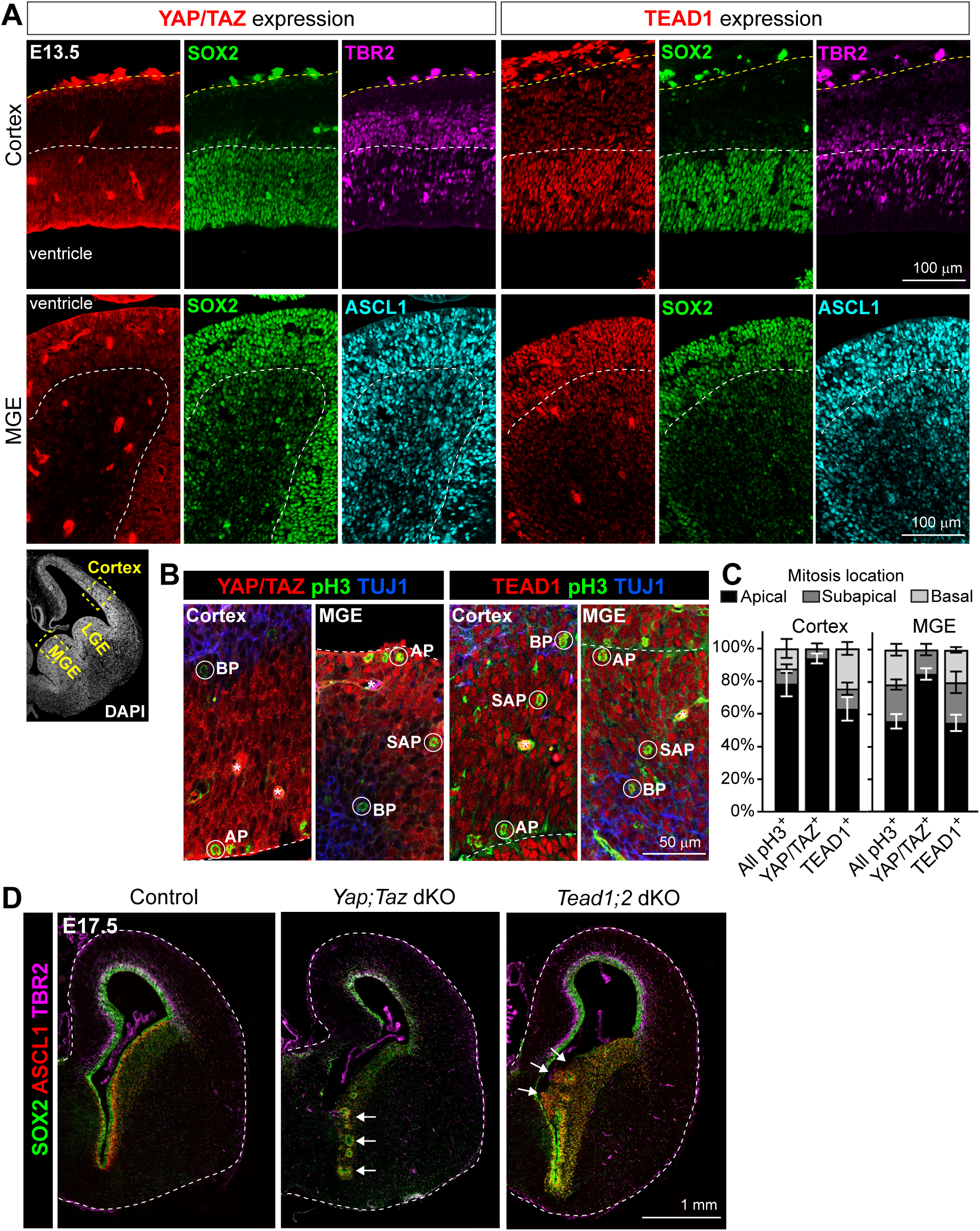
Expression patterns of YAP/TAZ and TEAD in the developing forebrain and gross phenotypes of their mutants. (A) YAP/TAZ and TEAD1 expression patterns in E13.5 forebrain sections examined by immunostaining. White dashed lines delineate the boundary between the ventricular zone (VZ) and the subventricular zone (SVZ). Yellow dashed lines mark the outer surface of the cortex. MGE, medial ganglionic eminence; LGE, lateral ganglionic eminence. (B) YAP/TAZ and TEAD1 immunostaining on E13.5 forebrain sections co-stained with phospho-histone H3 (pH3, cells in mitosis) and βIII-tubulin (TUJ1, neurons), highlighting apical progenitors (APs) that divide at the ventricular surface (VS), subapical progenitors (SAPs) that divided at subapical positions within the VZ, and basal progenitors (BPs) that divide in the SVZ. Dashed lines delineate the VS. Asterisks, autofluorescent blood vessels. (C) Quantification of the proportion of pH3^+^ cells at the respective locations. Values are mean ± SEM. (D) Immunostaining showing the gross phenotypes of *Nestin-Cre* mediated *Yap;Taz* double knockout (dKO) and *Tead1;2* dKO forebrain at E17.5. Dashed lines mark the outer surface of the forebrain. Arrows highlight disruptions in the VS and VZ-SVZ organization.

To further determine whether YAP/TAZ and TEAD1 are expressed in APs or BPs, we co-immunostained YAP/TAZ or TEAD1 with the mitosis marker phospho-Histone H3 (pH3) (Fig. 1B), because mitosis position is a defining feature of APs and BPs. In the cortex, >90% of YAP/TAZ^+^pH3^+^ cells were found along the VS (apical) and a small fraction within the VZ away from the VS (subapical) (Fig. 1C); the latter type of progenitors has been defined as SAPs— they are derived from APs and give rise to BPs (Pilz et al. 2013). Among TEAD1^+^pH3^+^ cells, only ∼60% were found along the VS, with the rest either within the VZ or in the SVZ (basal) (Fig. 1C). Similar patterns were observed in the MGE, except that the fraction of those dividing apically was reduced by ∼10% for both YAP/TAZ^+^ cells and TEAD1^+^ cells (Fig. 1C). These results demonstrate that, in the developing telencephalon, YAP/TAZ are predominantly expressed in RG/APs, whereas TEAD1 is expressed in both RG and IPs (including SAPs and BPs).

### YAP/TAZ loss and TEAD1/2 loss result in different defects in the subpallium

The above finding led us to hypothesize that TEAD may act independent of YAP/TAZ in IPs during telencephalon development. To test this, we decided to compare the phenotypes of genetic mutants lacking *Yap* and *Taz* (*Yap;Taz*) to those lacking *Tead*. Besides *Tead1*, *Tead2* mRNA is also detected in the VZ/SVZ throughout the developing CNS (Milewski et al. 2004) (Supplemental Fig. S1A). *Tead3* transcript levels in the telencephalon are very low (Supplemental Fig. S1A). Our RNA-sequencing (RNA-seq) data suggest that *Tead4* is not expressed in the developing CNS (data not shown). Therefore, we generated *Yap;Taz* double conditional knockout (dKO), *Tead1;2* dKO, and *Tead1* and *Tead2* single conditional KO (cKO) mice using a *Nestin-Cre* (*Nes-Cre*) line, which is expressed in the developing CNS starting from around embryonic day (E) 10.5. The depletion of corresponding proteins was validated by Western blot (Supplemental Fig. S1B).

Both *Yap;Taz* dKO and *Tead1;2* dKO mice died soon after birth. Therefore, we collected E17.5 brains for histological analysis. Obvious defects were observed in both mutants. In the subpallium, both mutants showed structural disruption of the VS and disorganization of the VZ-SVZ (Fig. 1D, arrows); similar defects were also present in the cortex of *Yap;Taz* dKO mice when *Yap;Taz* were deleted with *Emx1-Cre* (Lavado et al. 2021). Interestingly, *Tead1;2* dKO, but not *Yap;Taz* dKO, mice showed a marked expansion of ASCL1^+^ cells (Fig. 1D), suggesting that TEAD1/2 have YAP/TAZ-independent functions during subpallial development. Cortical neural progenitors, labeled by SOX2 and TBR2, were notably reduced in *Yap;Taz* dKO mice (Supplemental Fig. S1C), consistent with our previous report when *Yap;Taz* were deleted with *Emx1-Cre* (Lavado et al. 2021). This phenotype, however, was not present in *Tead1;2* dKO mice, suggesting either that TEAD1/2 are not required in cortical progenitors or that the loss of *Tead1;2* is compensated by other genes. Here we focused on investigating the role of TEAD1/2 during subpallial development.

The telencephalon of *Tead1* and *Tead2* single cKO mice did not show obvious defects at E17.5 (Supplemental Fig. S1D). However, *Tead1* cKO mice exhibited hydrocephaly postnatally (Supplemental Fig. S1D) and all died within 3 weeks of age. *Tead2* cKO mice did not show gross defects in brain development and were healthy and fertile. These results suggest that, functionally, *Tead1* and *Tead2* are largely redundant during brain development, although *Tead1* is probably more important than *Tead2*.

### TEAD1/2 loss and YAP/TAZ loss similarly disrupt subpallial apical junctions

The disruptions in the subpallial VS of E17.5 *Tead1;2* dKO and *Yap;Taz* dKO mice suggested defects in the apical junctions that support neuroepithelial structural integrity. Indeed, immunostaining using apical junction markers ZO-1, β-catenin, and aPKC revealed disruptions in these junctions as early as E13.5 in both dKO lines (Supplemental Fig. S2), indicating that TEAD1/2 and YAP/TAZ are both required to maintain the integrity of neuroepithelial apical junctions.

### TEAD1/2 loss and YAP/TAZ loss have distinct impacts on lineage progression of subpallial neural progenitors

To pinpoint which progenitor subtypes/states were affected upon TEAD1/2 loss and YAP/TAZ loss, we focused our analyses on the MGE. Previous studies have shown that nearly all cells in the VZ of MGE express OLIG2 as early as E10.5 (Petryniak et al. 2007), and that ASCL is expressed in cells both in the VZ and the SVZ (Yun et al. 2002; Petryniak et al. 2007; Imayoshi et al. 2013a). Studies have also found that the manner in which ASCL1 is expressed has distinct impact on cell fate; oscillatory expression maintains progenitor proliferation and self-renewal, whereas sustained expression promotes neuronal commitment and differentiation (Imayoshi et al. 2013b). We performed OLIG2, ASCL1, and pH3 triple immunostaining to correlate OLIG2 and ASCL1 expression patterns with progenitor subtype (Fig. 2A).

**Figure 2.**
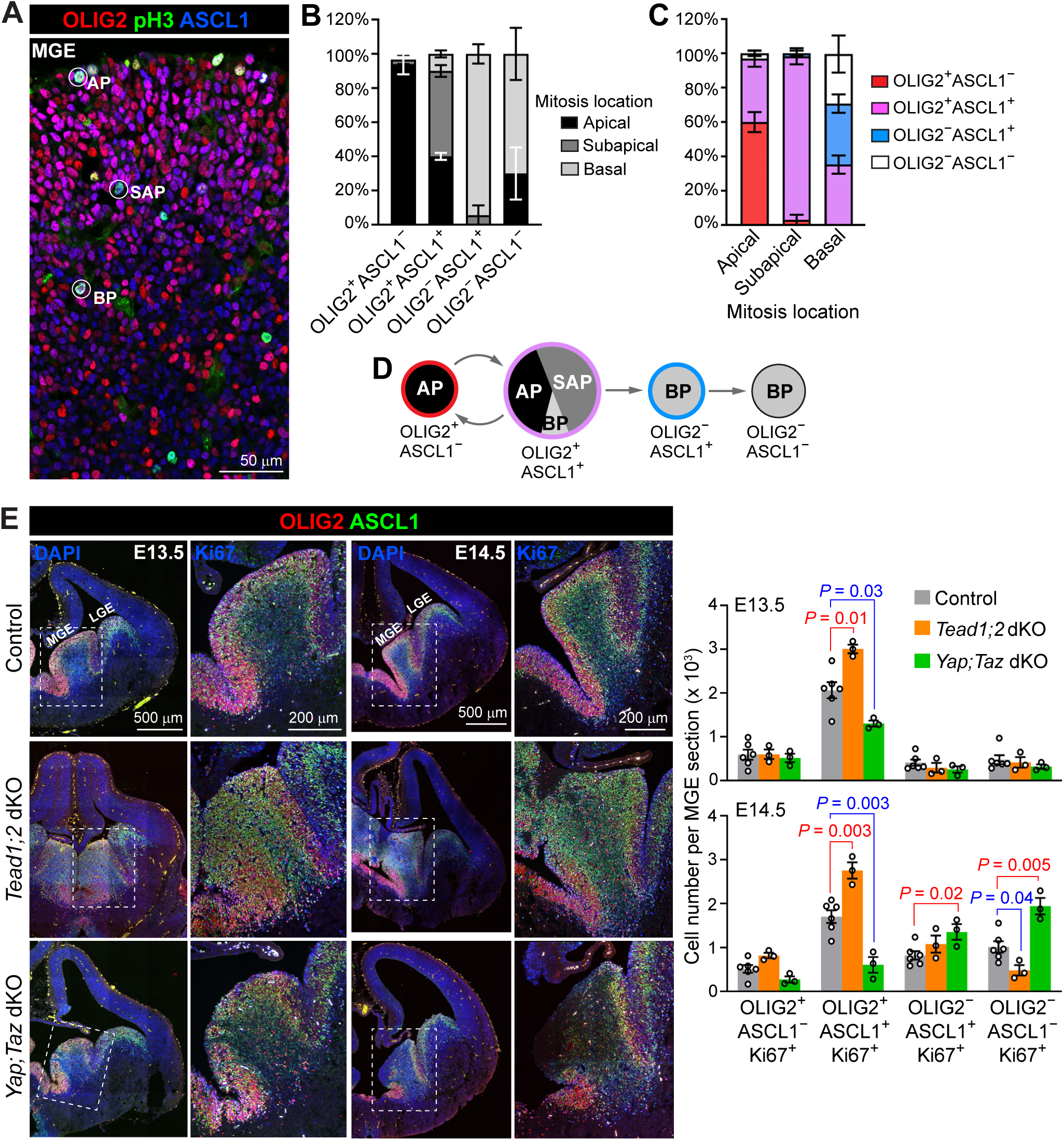
TEAD1/2 loss and YAP/TAZ loss have distinct impacts on subpallial neural progenitor lineage progression. (A) Immunostaining on a E13.5 brain section, highlighting an apical progenitor (AP), a subapical progenitor (SAP), and a basal progenitor (BP) in the MGE. (B and C) Quantification of the proportion of pH3^+^ cells found at different locations. (D) A schematic of proposed progenitor states during subpallial neural progenitor lineage progression. (E) Immunostaining and quantification of progenitors in the states defined by OLIG2 and ASCL1 immunosignal. N = 6 control mice with 3 from each line, N = 3 dKO mice. Areas in dashed boxes are enlarged in the images to the right. MGE, medial ganglionic eminence. LGE, lateral ganglionic eminence. Values are mean ± SEM. Each data point represents an individual animal. Two-sided unpaired *t*-test. Unmarked comparisons (vs. control) did not show significant difference (*P* > 0.05).

Quantification suggested that nearly all OLIG2^+^ASCL1^−^ cells were APs; that among OLIG2^+^ASCL1^+^ cells, ∼40% were APs, ∼50% were SAPs, and ∼10% were BPs; and that nearly all OLIG2^−^ASCL1^+^ and OLIG2^−^ASCL1^−^ progenitor cells were BPs (Fig. 2B). From another perspective: APs were composed of OLIG2^+^ASCL1^−^ (∼60%) and OLIG2^+^ASCL1^+^ (∼40%) cells; nearly all SAPs were OLIG2^+^ASCL1^+^ cells, and BPs were composed of roughly equal fractions of OLIG2^+^ASCL1^+^, OLIG2^−^ASCL1^+^, and OLIG2^−^ASCL1^−^ (and presumably Ki67^+^) cells (Fig. 2C). Taken together, we propose that MGE neural progenitors oscillate between the OLIG2^+^ASCL1^−^ and OLIG2^+^ASCL1^+^ states, then—upon neuronal commitment—progress from the OLIG2^+^ASCL1^+^ state to the OLIG2^−^ASCL1^+^ state, and finally to the OLIG2^−^ASCL1^−^ state (Fig. 2D). Although defining progenitor states in this manner is not ideal, as the OLIG2^+^ASCL1^+^ state contains cells in different cell states and some are probably no different from cells in the OLIG2^+^ASCL1^−^ state, this approach at least allows us to distinguish early progenitors (OLIG2^+^ASCL1^−^ and OLIG2^+^ASCL1^+^) and late progenitors (OLIG2^−^ASCL1^+^ and OLIG2^−^ASCL1^−^).

Next, we quantified the numbers of progenitors in each of the states defined above; the proliferation marker Ki67 was used as a generic marker for progenitor cells (Fig. 2E). In *Tead1;2* dKO MGE, OLIG2^+^ASCL1^+^ progenitors were significantly increased at E13.5 and E14.5 compared to control MGE while OLIG2^−^ASCL1^−^ progenitors significantly decreased at E14.5. In contrast, in *Yap;Taz* dKO MGE, OLIG2^+^ASCL1^+^ progenitors were significantly decreased at E13.5 and E14.5 while OLIG2^−^ASCL1^+^ and OLIG2^−^ASCL1^−^ progenitors significantly increased at E14.5. In both dKO lines, the numbers of OLIG2^+^ASCL1^−^ cells were not significantly changed (Fig. 2E). Thus, TEAD1/2 loss and YAP/TAZ loss have distinct impacts on the lineage progression of subpallial progenitors: TEAD1/2 loss expands the early progenitor (OLIG2^+^ASCL1^+^) population, whereas YAP/TAZ loss depletes early progenitors and expands late (OLIG2^−^) progenitors.

### TEAD1/2 loss and YAP/TAZ loss differentially affect MGE cellular output

MGE progenitors give rise to cortical interneurons, projection neurons and interneurons in the basal ganglia, and OLCs in the embryonic cortex (Kessaris et al. 2006; Rubenstein and Campbell 2013; Mayer et al. 2018). In *Tead1;2* dKO mice, the percentage of TUJ1^+^ area within the MGE and the number of MAFB^+^ immature interneurons (Pai et al. 2019) in the neocortex were significantly reduced at E14.5 compared to those in control mice (Fig. 3A and 3B). Cortical OLCs labeled by OLIG2 and SOX10 (Rowitch 2004) were also significantly reduced in *Tead1;2* dKO mice at E17.5 (Fig. 3C and 3D). These results suggest that TEAD1/2 loss impairs the production of neurons and OLCs by the MGE.

**Figure 3.**
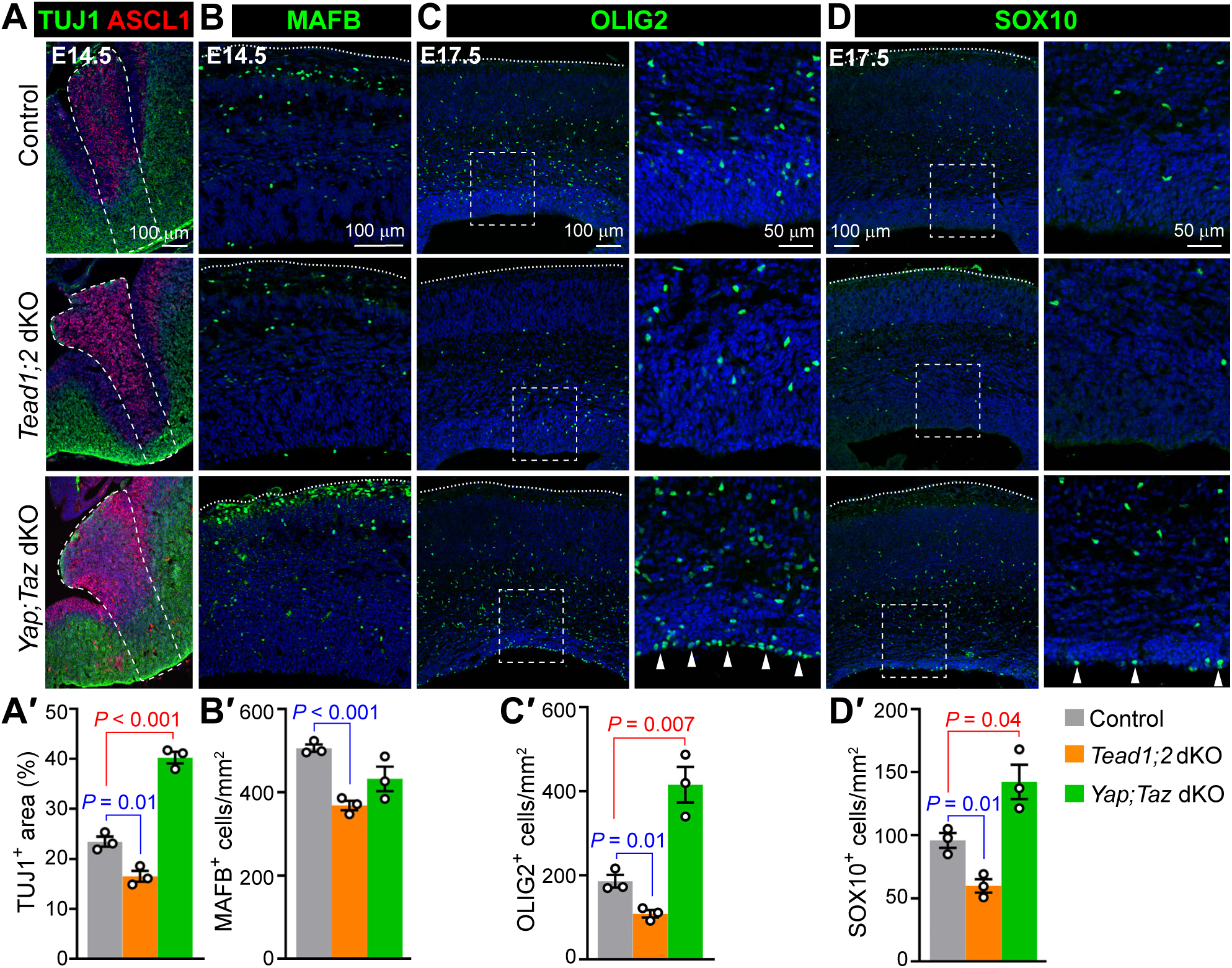
TEAD1/2 loss and YAP/TAZ loss have distinct impacts on the cellular output from subpallial progenitors. (A) Immunostaining and quantification of neurons (TUJ1) in the MGE (dashed area). (B) Immunostaining and quantification of immature interneurons (MAFB) in the neocortex. (C and D) Immunostaining and quantification of oligodendrocyte lineage cells (OLIG2 and SOX10) in the neocortex. Areas in dashed boxes are enlarged in the images to the right. Arrowheads highlight OLIG2- and SOX10-labeled cells located at the ventricular surface. Blue color is DAPI signal. Dotted lines mark the outer surface of the cortex. Values are mean ± SEM. Each data point represents an individual animal. Two-sided unpaired *t*-test. Unmarked comparisons (vs. control) did not show significant difference (*P* > 0.05).

In *Yap;Taz* dKO mice, the percentage of TUJ1^+^ area within the MGE was increased at E14.5 compared to control mice, although the number of MAFB^+^ immature interneurons in the neocortex was unchanged (Fig. 3A and 3B). Cortical OLCs were increased in *Yap;Taz* dKO mice at E17.5; intriguingly, many OLIG2- and SOX10-labeled cells were found along cortical VS in *Yap;Taz* dKO (Fig. 3C and 3D, arrowheads), which were rarely seen in control mice. These results suggest that YAP/TAZ loss causes premature neuronal differentiation. Whether YAP/TAZ loss enhances OLC differentiation by MGE progenitors remains to be determined because it is unclear whether the excess OLCs found *Yap;Taz* dKO cortices were derived from MGE or cortical progenitors. Taken together, our results demonstrate that TEAD1/2 loss and YAP/TAZ loss differentially affect the cellular output from MGE neural progenitors.

### Single-cell RNA-seq analysis reveals distinct impacts of TEAD1/2 loss and YAP/TAZ loss on MGE neural progenitor lineage progression

To confirm the changes in MGE progenitor population in our mutant mice and to hopefully better define progenitor states, we performed single-cell RNA-seq (scRNA-seq) using microdissected MGE from E14.5 *Tead1;2* dKO, *Yap;Taz* dKO, and their corresponding no-*Cre* littermate controls, as well as MGE from E12.5 and E15.5 wild-type (WT) mice.

To understand MGE lineage progression during normal development, we first analyzed WT cells, which included E12.5 and E15.5 WT cells and E14.5 *Tead1;2* control cells. We clustered them based on highly variable genes (HVGs) and visualized them by uniform manifold approximation and projection (UMAP) (Fig. 4A and Supplemental Fig. S3A). To annotate the clusters, we compiled a list of progenitor genes and a list of neuron genes (Supplemental Table S1) based on previous publications (Mayer et al. 2018; Mi et al. 2018) and scored each cell based on their expression of these genes. This analysis indicated that clusters 2, 3, and 6 were predominantly composed of progenitor cells and clusters 1, 4, 5, 7, and 8 were composed of neurons (Fig. 4A–C). This conclusion is supported by cell-cycle phase analysis, which showed that cells in clusters 2, 3, and 6 were predominantly proliferating cells in S and G2/M phases, and cells in clusters 1, 4, 5, 7, and 8 were postmitotic cells in G1/G0 phases (Fig. 4D). Based on marker gene expression, cells in cluster 9 were endothelial cells (*Igfbp7*), in cluster 10 were red blood cells (*hba* and *hbb*), and in cluster 11 were microglia (*Cx3cr1*).

**Figure 4.**
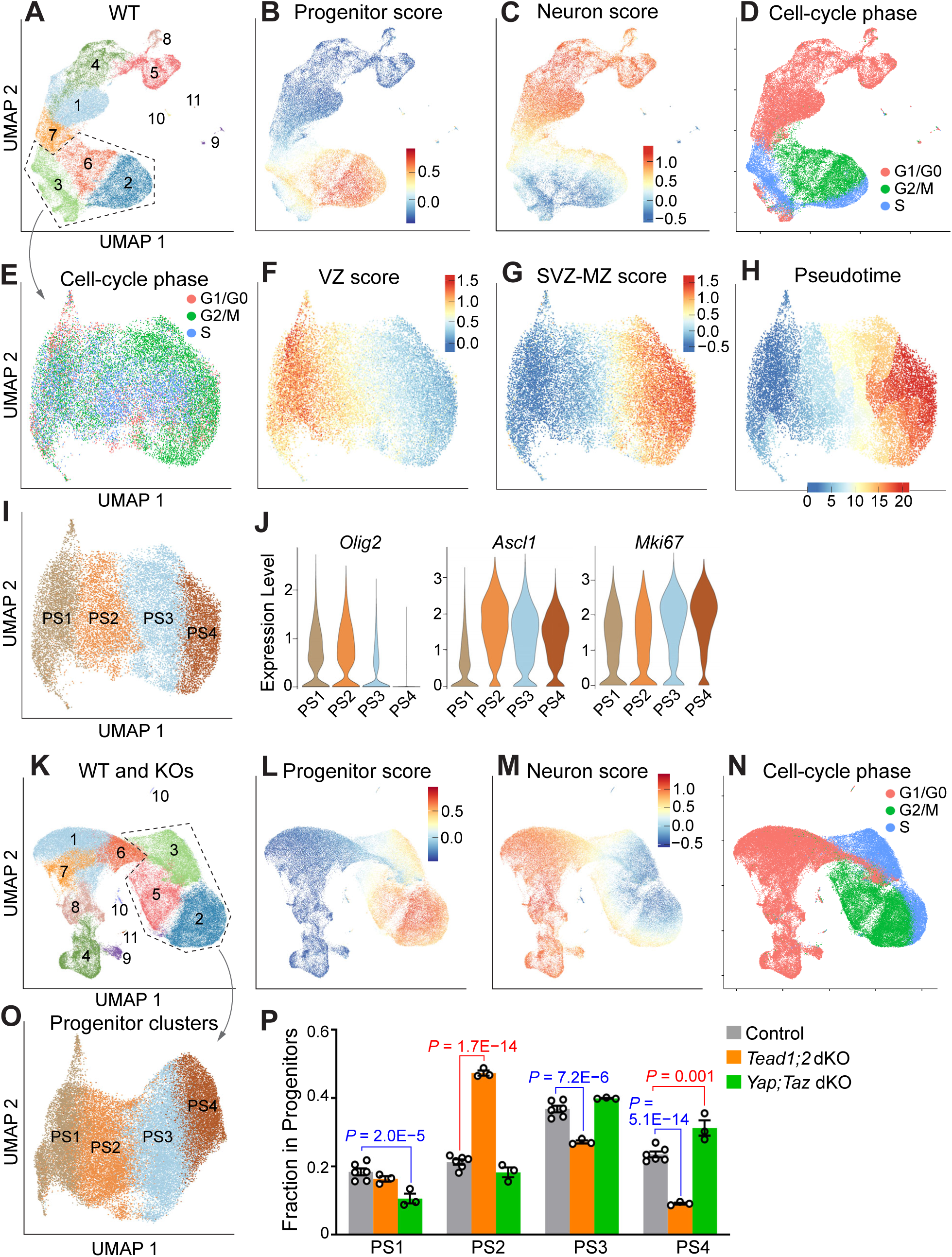
Single-cell RNA sequencing analysis reveals distinct impacts of TEAD1/2 loss and YAP/TAZ loss on MGE neural progenitor lineage progression. (A) Clustering of WT MGE cells (which included E12.5 WT, E14.5 *Tead1;2* control, and E15.5 WT cells; n = 42,876 cells after quality control) based on highly variable genes (HVG) visualized by UMAP. (B–D) UMAP visualizations of progenitor score, neuron score, and cell-cycle phase of WT MGE cells. (E) Cell cycle phase of WT progenitor cells (cells from clusters 2, 3, and 6 in A; n = 18,963 cells after quality control) visualized by UMAP using a curated list of transcription factors for feature selection. (F–H) UMAP visualizations of VZ score, SVZ-MZ score, and pseudotime of WT progenitor cells. (I) Clustering of WT progenitor cells based on the list of transcription factors at the resolution of 0.25. PS, progenitor state. (J) ScRNA-seq expression levels of indicated genes in each progenitor state. (K) Clustering of WT and KO MGE cells (which included E12.5 WT, E14.5 *Tead1;2* control and dKO, E14.5 *Yap;Taz* control and dKO, and E15.5 WT cells; n = 85,684 cells after quality control) based on HVG visualized by UMAP. (L–N) UMAP visualizations of progenitor score, neuron score, and cell-cycle phase of WT and KO cells. (O) Progenitor state annotation of WT and KO MGE progenitor cells (cells from clusters 2, 3, and 5 in K; n = 38,860 cells after quality control) visualized by UMAP using the list of transcription factors for feature selection. (P) Fraction of progenitor cells in each progenitor state. Values are mean ± SEM. Each data point represents an individual animal. Statistical test was performed using *propeller*. Unmarked comparisons (vs. control) did not show significant difference (*P* > 0.05).

Next, we subset out the progenitor clusters (clusters 2, 3, and 6) for further analysis. When HVGs were used for feature selection, progenitor cells were organized largely based on their cell-cycle phase; for example, cluster 3 was mostly composed of S-phase cells and cluster 6 was composed of G2/M-phase cells (Fig. 4D). A previous study of subpallial development has found that clustering cells using transcription factors mitigates the influence of cell-cycle phase and improves resolution of developmental populations (Su-Feher et al. 2022). We adopted this approach and compiled a list of 742 transcription factor genes (Supplemental Table S1) based on both the aforementioned study and the Allen Developing Mouse Brain Atlas RNA in situ hybridization data. Using this list for feature selection, cells in different cell-cycle phases intermingled when visualized by UMAP (Fig. 4E), indicating that the influence of cell-cycle phase was largely abolished. Scoring the cells using our curated lists of genes enriched in the VZ and in the SVZ and mantle zone (SVZ-MZ) (Supplemental Table S1 and Supplemental Fig. S3B) showed a gradual, unidirectional transition of VZ and SVZ-MZ scores on UMAP (Fig. 4F and 4G). Pseudotime analysis by Monocle 3 also revealed a gradual, unidirectional transition (Fig. 4H). Together, these results suggest that using our list of transcription factors to cluster MGE progenitor cells is able to capture their lineage progression.

How many MGE progenitor states/subtypes are there? To address this question, we used an unbiased, systematic approach by stepwise (0.025) increasing the clustering resolution and plotting the results in a “cluster tree” (Supplemental Fig. S3C). When the resolution was less than 0.25, cells often changed their cluster membership at the next resolution (Supplemental Fig. S3C, grey diagonal arrows), suggesting that the clusters were unstable. When the resolution was 0.25 and higher, however, cells no longer changed their cluster membership other than splitting into smaller clusters (Supplemental Fig. S3C). We therefore concluded that 0.25 was the optimal resolution, yielding 4 clusters with comparable size, which we referred to as progenitor state (PS) 1 to PS4 with PS1 being the earliest progenitor state and PS4 being the latest (Fig. 4I). Expression counts of *Olig2* were high in PS1 and PS2 and low in PS3 and PS4, whereas that of *Ascl1* were low in PS1, high in PS2 and PS3, and medium in PS4 (Fig. 4J). These results match well with the progenitor states defined by OLIG2 and ASCL1 immunostaining (Fig. 2D), validating our computational methods to analyze and annotate MGE progenitor cells.

Next, we combined WT, *Tead1;2* dKO, *Yap;Taz* dKO, and no-*Cre* controls cells, clustered them based on HVGs (Fig. 4K and Supplemental Fig. S3D), and identified progenitor clusters— clusters 2, 3, and 5—based on progenitor/neuron scoring and cell-cycle phase analysis (Fig. 4L–N). We then assigned progenitor states by using the annotation of WT progenitor cells as the reference (Stuart et al. 2019) (Fig. 4O and Supplemental Fig. S3E). To compare the distribution of progenitor cells in each progenitor state between control and dKO samples, we used *propeller*, a method for testing for differences in cell type proportions in scRNA-seq data (Phipson et al. 2022). In *Tead1;2* dKO samples, the proportion of progenitor cells in PS2 was significantly increased and the proportions in PS3 and PS4 significantly decreased compared to control. In contrast, in *Yap;Taz* dKO samples, the proportion of progenitor cells in PS1 was significantly decreased and the proportion in PS4 significantly increased (Fig. 4P). These results correlate well with that obtained by immunostaining (Fig. 2E); both approaches find that TEAD1/2 loss expands the early progenitor population and depletes later progenitors, whereas YAP/TAZ loss has the opposite effect—it depletes early progenitors and expands late progenitors. Pseudotime analysis further supports this conclusion (Supplemental Fig. S3F).

### TEAD1/2 loss impedes developmental progression of subpallial neural progenitors

The consequence of YAP/TAZ loss in the subpallium, a depletion of early progenitors, is consistent with the known function of YAP/TAZ in promoting the proliferation and self-renewal of diverse types of stem/progenitor cells (Driskill and Pan 2023; Zhong et al. 2024). However, that TEAD1/2 loss results in an expansion of early progenitors is unexpected. Therefore, we investigated the cellular mechanism driving this expansion.

A faster cell-cycle speed could lead to increased cell numbers within a given time window. We measured cell-cycle speed using a sequential EdU-BrdU labeling method (Quinn et al. 2007). The cell-cycle length and the S-phase length of OLIG2^+^ASCL1^+^ cells—the population expanded in *Tead1;2* dKO MGE—and of OLIG2^+^ASCL1^−^ cells were significantly increased in *Tead1;2* dKO MGE compared to control (Fig. 5A), indicating a slowing down of cycling speed. This could not explain why the number of OLIG2^+^ASCL1^+^ cells was increased in *Tead1;2* dKO MGE.

**Figure 5.**
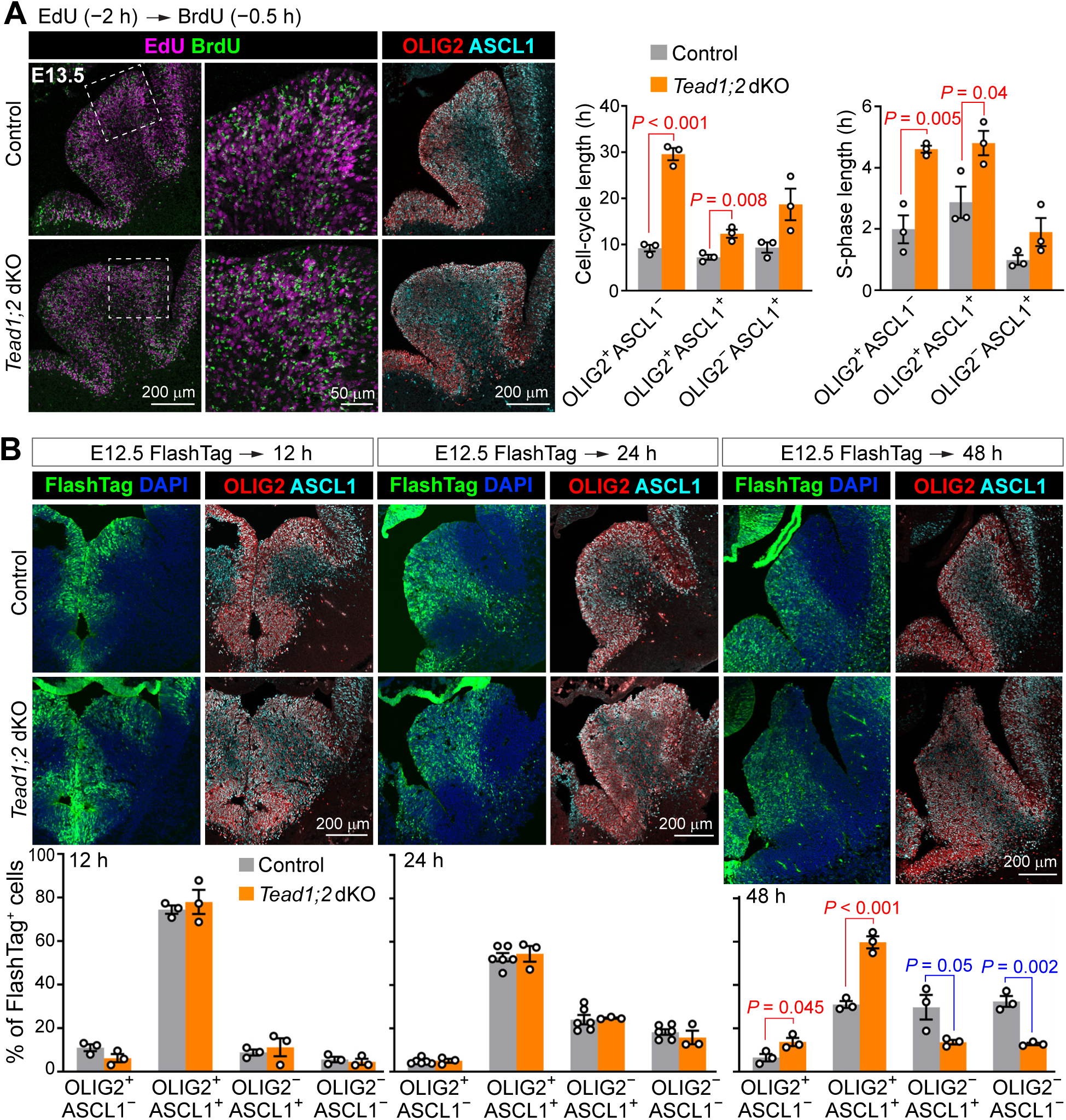
TEAD1/2 loss impedes developmental progression of subpallial neural progenitors. (A) Immunostaining and quantification measuring the cell cycle speed of MGE progenitor cells by EdU-BrdU sequential labeling. Areas in dashed boxes are enlarged in the images to the right. (B) Immunostaining and quantification of the fraction of FlashTag-labeled cells in the progenitor states defined by OLIG2 and ASCL1 immunosignal. Values are mean ± SEM. Each data point represents an individual animal. Two-sided unpaired *t*-test. Unmarked comparisons (vs. control) did not show significant difference (*P* > 0.05).

Next, we tested whether the developmental progression of early progenitors was stalled in *Tead1;2* dKO MGE. To this end, we performed lineage tracing by injecting FlashTag into the cerebral ventricles of E12.5 embryos in utero to pulse label M-phase APs(Govindan et al. 2018) and monitoring the fate of labeled cells 12, 24, and 48 h post injection (hpi). At 12 and 24 hpi, the fate of FlashTag-labeled cells in control and *Tead1;2* dKO MGE was similar (Fig. 5B); in both genotypes the fractions of labeled cells existing in early progenitor (OLIG2^+^) states decreased while those adopting later progenitor (OLIG2^−^) states increased comparing 24 hpi vs. 12 hpi (Fig. 5B), consistent with developmental progression. At 48 hpi, however, significantly larger fractions of labeled cells remained in the OLIG2^+^ early progenitor states—especially the OLIG2^+^ASCL1^+^ state—rather than progressing to the OLIG2^−^ late progenitor states in *Tead1;2* dKO than in control MGE (Fig. 5B). Thus, TEAD1/2 promote developmental progression of subpallial neural progenitors; their loss stalls the cells in early progenitor states.

### TEAD1/2 promote lineage progression of subpallial progenitors by inhibiting Notch signaling

To uncover the molecular mechanism underlying the expansion of early progenitors in *Tead1;2* dKO MGE, we exploited our scRNA-seq dataset and performed differential expression analysis using scMINER (Ding et al. 2023) and gene set enrichment analysis (GSEA) using NetBID2 (Dong et al. 2023), comparing *Tead1;2* dKO and control PS2 cells, the progenitor state that was expanded in *Tead1;2* dKO MGE (Supplemental Table S2). Several signaling pathway gene sets in the Hallmark collection were significantly upregulated in *Tead1;2* dKO PS2 cells, among which Notch_signaling was upregulated the most strongly (Fig. 6A and 6B). Interestingly, several cell cycle related gene sets, such as E2F_targets and G2M_checkpoint, were significantly downregulated in *Tead1;2* dKO PS2 cells (Fig. 6A and Supplemental Table S2), which is consistent with our finding that the cell-cycle speed of early progenitor cells was slowed down in *Tead1;2* dKO MEG.

**Figure 6.**
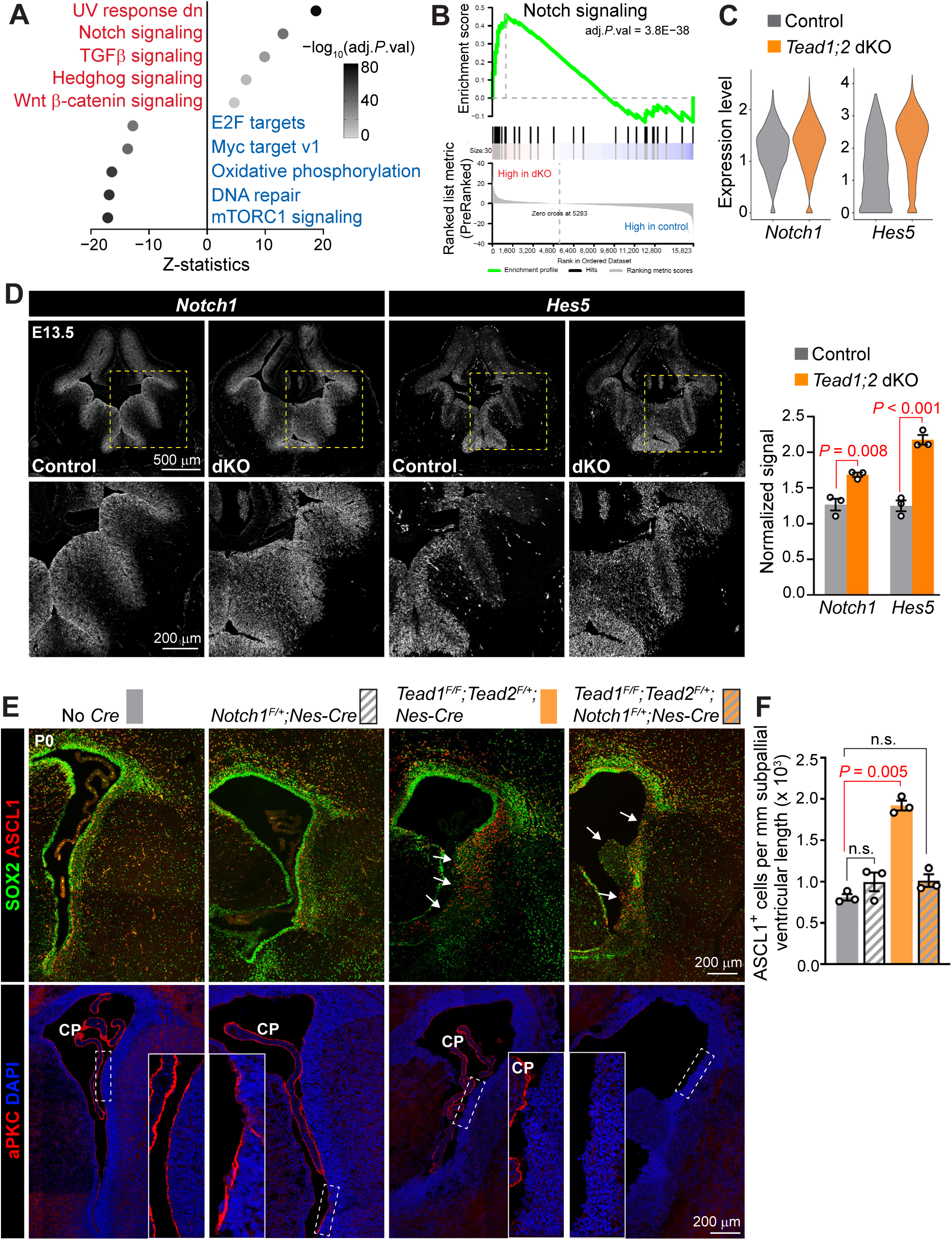
TEAD1/2 promote lineage progression of subpallial neural progenitors by inhibiting Notch signaling. (A) Top 5 enriched Hallmark gene sets by gene set enrichment analysis (GSEA) comparing *Tead1;2* dKO to control PS2 cells. (B) Enrichment plot of the Notch_signaling gene set. (C) ScRNA-seq expression levels of *Notch1* (log_2_FC = 0.17, adj.*P*.val = 0.001) and *Hes5* (log_2_FC = 2.24, adj.*P*.val = 2.8E−143) in PS2 cells comparing *Tead1;2* dKO vs. control. (D) RNAscope in situ hybridization analysis of *Notch1* and *Hes5* expression in E13.5 brain sections. Areas in dashed boxes are enlarged in the images below. Mean fluorescence intensity in the MGE (total signals divided by area) were normalized to the mean intensity in a cortical VZ region within the same section. (E) Immunostaining of neural progenitors (SOX2 and ASCL1) and apical junctions (aPKC) in P0 brain sections. Arrows highlight disruptions in the VS and VZ-SVZ organization. CP, choroid plexus. (F) Quantification of ASCL1^+^ cells located in subpallial VZ-SVZ in P0 brain sections. Values in D and F are mean ± SEM. Two-sided unpaired *t*-test; n.s., not significant (*P* > 0.05). Each data point represents an individual animal.

Notch signaling is a key regulator of neural progenitor cells. Conditional deletion of *Notch1* leads to accumulation of *Ascl1*-expressing cells in the subpallial VZ while depleting these cells in the SVZ (Mason et al. 2005). *Notch1–3* and Notch target genes, *Hes1* and *Hes5*, were all significantly upregulated in *Tead1;2* dKO PS2 cells compared to control PS2 cells (Fig. 6C and Supplemental Table S2). We confirmed the upregulation of *Notch1* and *Hes5*—the most highly expressed *Notch* and *Hes* genes in PS2 cells—by RNAscope (Fig. 6D).

To determine whether upregulation of Notch signaling contributes to the expansion of early progenitors in *Tead1;2* dKO mice, we performed genetic rescue experiments by deleting one allele of *Notch1* in the *Tead1;2* dKO background. This, unfortunately, did not rescue the subpallial phenotype of *Tead1;2* dKO mice (data not shown). We reasoned that the failure of rescue was probably because the phenotype of *Tead1;2* dKO mice was too severe. Therefore, we attempted the rescue in the *Tead1^F/F^;Tead2^F/+^;Nes-Cre* background; these mice also exhibited consistent expansion of the ASCL1^+^ progenitor population at postnatal day (P) 0 (Fig. 6E and 6F), although not as severely as *Tead1;2* dKO mice did. Loss of one allele of *Notch1*, which had no effect on the number of ASCL1^+^ cells in the WT background, completely restored the number of ASCL1^+^ cells in *Tead1^F/F^;Tead2^F/+^;Nes-Cre* mice to the control level (Fig. 6E and 6F). These results indicate that TEAD1/2 promote lineage progression of subpallial neural progenitors at least in part by inhibiting Notch signaling. Note that in *Tead1^F/F^;Tead2^F/+^;Nes-Cre* subpallium, the VS, apical junctions, and VZ-SVZ organization were disrupted (Fig. 6E, arrows and insets), just like in *Tead1;2* dKO mice. These defects, however, were not rescued by the loss of one allele of *Notch1* (Fig. 6E, arrows and insets), suggesting that upregulation of Notch signaling is not responsible for these defects.

### TEAD1/2 act with INSM1 to promote lineage progression of subpallial neural progenitors

The distinct subpallial phenotypes of *Tead1;2* dKO and *Yap;Taz* dKO mice strongly suggest that TEAD1/2 regulate lineage progression of subpallial neural progenitors independent of YAP/TAZ. In addition to Yorki/YAP/TAZ, Scalloped/TEAD interacts with Vestigial/VGLL1–3 (Halder et al. 1998; Simmonds et al. 1998), Tgi/VGLL4 (Guo et al. 2013; Koontz et al. 2013), Nerfin-1/INSM1 (Vissers et al. 2018; Guo et al. 2019), and RUNX2/3 (Qiao et al. 2016; Suo et al. 2020), among which only *Vgll4* (Vestigial like 4) and *Insm1* are expressed in subpallial progenitor cells (Supplemental Fig. S4A). By competing with YAP/TAZ for TEAD binding, VGLL4-mediated transcriptional repression antagonizes YAP/TAZ-mediated transcriptional activation of TEAD target genes (Lin et al. 2016; Cai et al. 2022). To test whether TEAD1/2 act with VGLL4 to regulate subpallial lineage progression, we generated *Vgll4* null mice and *Nes-Cre*–mediated cKO mice. Null mice all died before P7 (Supplemental Fig. S4B), an outcome reported previously (Yu et al. 2019). However, cKO mice did not exhibit obvious brain development phenotype (Supplemental Fig. S4D), suggesting that VGLL4 is not an important cofactor for TEAD in the context of subpallial development.

INSM1 is a zinc-finger transcription factor with an N-terminal SNAG (Snail/Gfi-1) domain that interacts with transcriptional corepressors such as KDM1A/LSD1 (lysine (K)-specific demethylase 1A) (Welcker et al. 2013). *Insm1* mRNA is expressed in neural progenitors and nascent neurons throughout embryonic and adult neurogenesis (Duggan et al. 2008). Within neural progenitors, *Insm1* is expressed in non-surface dividing SAPs and BPs but not in APs (Duggan et al. 2008; Farkas et al. 2008; Tavano et al. 2018). Previous studies have found that *Insm1* deletion leads to increased subpallial VZ/SVZ size and the number of Ki67^+^ cells and decreased embryonic cortical OLCs (Farkas et al. 2008; Monaghan et al. 2017)—similar to the effects of *Tead1;2* deletion, raising the possibility that TEAD1/2 act with INSM1 to regulate subpallial development. To test this, we examined INSM1 expression pattern in the MGE (Fig. 7A). At E12.5, INSM1 was expressed in some VZ cells and more frequently in SVZ cells. All INSM1^+^ cells in the VZ and a subset of INSM1^+^ cells in the SVZ, especially those close to the VZ, also expressed TEAD1.

**Figure 7.**
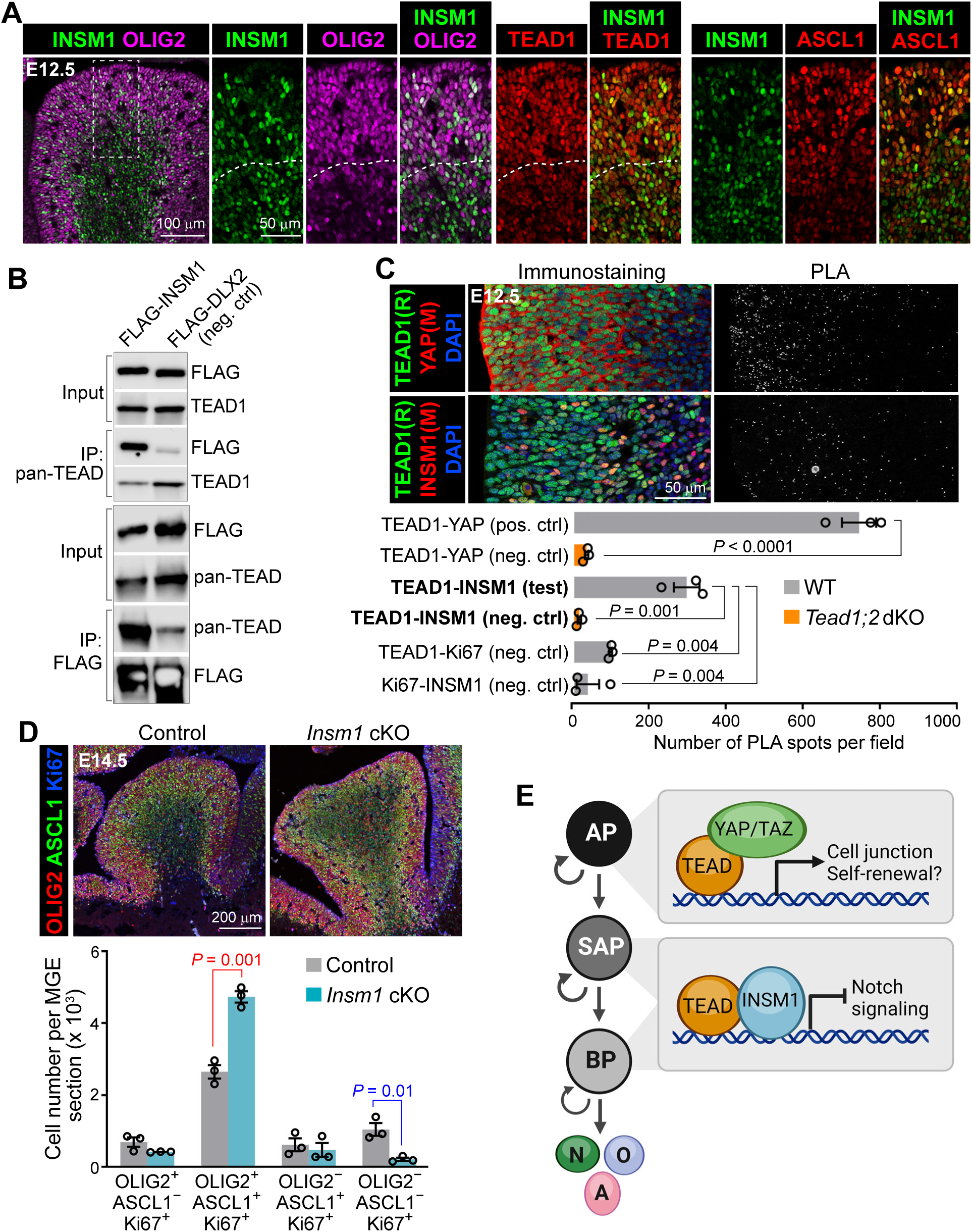
INSM1 interacts with TEAD and promote lineage progression of subpallial neural progenitors. (A) INSM1 expression pattern in E12.5 MGE examined by immunostaining. Dashed lines delineate VZ and SVZ boundary. (B) Co-immunoprecipitation experiment using 293T cells transfected with FLAG-tagged INSM1 or DLX2 (negative control) to examine the interaction between endogenous TEAD and FLAG-tagged proteins. (C) Immunostaining and proximity ligation assay (PLA) images of adjacent E12.5 brain sections showing subpallial regions and quantification of PLA signals as fluorescent spots. R, rabbit antibody; M, mouse antibody. (D) Immunostaining and quantification of neural progenitors in E14.5 MGE. Values are mean ± SEM. Each data point represents an individual animal. Two-sided unpaired *t*-test. Unmarked comparisons (vs. control) did not show significant difference (*P* > 0.05). (E) A model of TEAD function during subpallial development. TEAD interacts with different partners in different progenitor subtypes/states to execute distinct functions. In apical progenitors (APs), TEAD interacts with YAP/TAZ to promote the expression of cell junction genes; whether TEAD also promotes self-renewal with YAP/TAZ remains to be determined. In subapical progenitors (SAPs) and possibly a subset of basal progenitors (BPs), TEAD interacts with INSM1 to promote lineage progression at least in part by inhibiting Notch signaling. N, neurons; O, oligodendrocytes; A, astrocytes. Created with BioRender.com.

We validated TEAD–INSM1 interaction by co-immunoprecipitation using transfected HEK293T cells (Fig. 7B). Proximity ligation assay (PLA) using E12.5 brain sections further confirmed TEAD–INSM1 interaction in the subpallium (Fig. 7C). Interestingly, whereas the PLA-detected interactions (PLA^+^ spots) between TEAD1 and YAP were enriched near the ventricular surface, the interactions between TEAD1 and INSM1 were more frequent near the VZ-SVZ boundary (Fig. 7C).

Previous characterizations of *Insm1* KO mice did not define the affected progenitor subtype/state with molecular markers. Therefore, we generated *Insm1* cKO mice using *Nes-Cre*. The loss of INSM1 protein was confirmed by Western blot (Supplemental Fig. S5A). At E14.5, the OLIG2^+^ASCL1^+^ progenitor population was significantly expanded and the OLIG2^−^ASCL1^−^ population significantly decreased in *Insm1* cKO mice compared to control mice (Fig. 7D). The subpallial ASCL1^+^ population remained expanded at P0 (Supplemental Fig. S5C). These phenotypes recapitulate that of *Tead1;2* dKO mice, supporting the hypothesis that TEAD1/2 and INSM1 act together to promote lineage progression of subpallial neural progenitors. Notably, the VS, apical junctions, and VZ-SVZ organization were not disrupted in *Insm1* cKO mice (Supplemental Fig. S5B), unlike in *Tead1;2* dKO mice; this is consistent with the idea that TEAD1/2 act with YAP/TAZ to maintain the integrity of neuroepithelial apical junctions.

## Discussion

In this study, we discovered that the TEAD family of transcription factors, best known as the DNA-binding factor of the Hippo pathway, execute distinct functions along ventral telencephalon lineage progression by interacting with different cofactors in different progenitor subtypes/states.

Prompted by the hypothesis that TEAD may have YAP/TAZ-independent functions during vertebrate neural tube development, we compared conditional mouse mutants lacking *Tead1* and *Tead2* (*Tead1;2* dKO) to those lacking *Yap* and *Taz* (*Yap;Taz* dKO). The distinct, essentially opposite, phenotype in the subpallium of these two mutant lines confirms our hypothesis. YAP/TAZ loss results in decreased numbers of subpallial early progenitors and increased numbers of late progenitors and neurons, indicative of reduced early progenitor self-renewal and premature differentiation, which is consistent with the well-established function of YAP/TAZ in promoting stem/progenitor cell self-renewal. In striking contrast, TEAD1/2 loss results in an expansion of early progenitors and reduction in late progenitors, neurons, and OLCs, suggesting that TEAD1/2 promote developmental progression of subpallial neural progenitors. One phenotype, however, is present in both mutant lines: the disruption of subpallial VS and VZ-SVZ organization, likely caused by defects in neuroepithelial apical junctions. We have shown that YAP/TAZ regulate the expression of cell junction genes in cortical neural progenitors (Lavado et al. 2021); the shared junctional defect suggests that YAP/TAZ and TEAD act together to regulate these genes. Taken together, our study demonstrates that, during mammalian subpallial development, TEAD has both YAP/TAZ-dependent and -independent functions.

Interestingly, despite having similar expression patterns in cortical and subpallial neural progenitors, TEAD1/2 loss specifically perturbs subpallial, but not cortical, development. It is unclear whether TEAD function is uniquely required in subpallial progenitors, or whether *Tead3* and/or *Tead4* compensate for the loss of *Tead1*;*2* in cortical progenitors. In comparison, YAP/TAZ loss affects the structural organization and self-renewal of cortical and subpallial progenitors similarly. Furthermore, in addition to having fewer early neural progenitors and more late progenitors and neurons in the MGE, *Yap;Taz* dKO embryos had more OLCs in the cortex. Interestingly, many OLCs were found along the VS in *Yap;Taz* dKO cortices, which rarely occurs in control embryos. Because cortical progenitors lacking YAP/TAZ are defective in their self-renewal and proliferation capacities (Lavado et al. 2021), it is possible that the OLCs located at VS were derived from cortical progenitors as a result of their abnormal differentiation, rather than from subpallial progenitors; resolving this will require future experiments using cortical and MGE region-specific *Yap;Taz* mutants.

Unlike in the cortex where RG/APs and IPs/BPs are readily distinguished by immunostaining (e.g., RG are SOX2^+^TBR2^−^ and IPs are TBR2^+^), progenitor subtype/state in the subpallium has not been defined in a similar manner. We used the combined immunosignals of OLIG2 and ASCL1—because their expression patterns are well characterized and because excellent antibodies are available—and defined four subpallial progenitor states. Although these progenitor states likely reflect the lineage progression of subpallial progenitors and, importantly, allowed us to distinguish early vs. late progenitors, there are several weaknesses with this approach. First, the four progenitor states do not progress in a linear fashion; cells in the OLIG2^+^ASCL1^−^ state oscillate with a subset of cells in the OLIG2^+^ASCL1^+^state due to the oscillatory expression of ASCL1 (Imayoshi et al. 2013b). Second, the OLIG2^+^ASCL1^+^ state contains several subtypes of progenitors (APs/RG, SAPs, and BPs). Lastly, this approach does not distinguish RG and IPs. Therefore, although our analysis shows that the OLIG2^+^ASCL1^+^ early progenitor population is affected—increased in *Tead1;2* dKO mice and decreased in *Yap;Taz* dKO mice, it remains unclear whether RG/APs, SAPs, or BPs are affected; using mitotic location to distinguish them is not feasible due to the disorganization of VZ-SVZ in both mutant lines.

To complement immunohistological analysis and overcome at least some of the weaknesses mentioned above, we employed scRNA-seq to characterize our mutant lines because this approach offers a more comprehensive view of cell state by measuring the expression levels of thousands of genes. Previous scRNA-seq analyses of the developing mouse subpallium did not define progenitor states in detail (Mayer et al. 2018; Mi et al. 2018; Allaway et al. 2021). By using a curated list of 742 transcription factors for feature selection and using stepwise increase of clustering resolution to identify the optimal cluster number, we defined four MGE progenitor states. Based on marker gene expression (*Olig2* and *Ascl1*), the four progenitor states defined by scRNA-seq—or more precisely, by the expression of the 742 transcription factors—correlate well with the four immunostaining-defined states. However, the progenitor states defined by these two methods are not identical. For example, in WT E14.5 MGE, the OLIG2^+^ASCL1^+^ state contains about twice the number of cells than the other states do, whereas the four progenitor states defined by scRNA-seq are of comparable size, with the third state (PS3) being the largest. More importantly, both gene signature analysis using VZ and SVZ-MZ gene scores and pseudotime analysis showed gradual, unidirectional transition of scRNA-defined progenitor states, rather than the oscillatory behavior between the two early progenitor states defined by immunostaining. Given that scRNA-based analysis takes into account much richer information and is much less biased than immunostaining-based analysis (which is limited by antibody availability and influenced by definition of immunosignal positivity), it is likely that the progenitor states defined by scRNA-seq reflect more accurately the true states and transition of progenitor cells during development. Our scRNA-seq analysis probably detected at least some of the different states of OLIG2^+^ASCL1^+^ cells—perhaps those in the self-renewal state (that oscillate with the OLIG2^+^ASCL1^−^ state) vs. those committed to differentiation—and separated them into two progenitor states, PS1 and PS2, respectively. Interestingly, our scRNA-seq analysis found that YAP/TAZ loss and TEAD1/2 loss affect different early progenitor states: YAP/TAZ loss reduces PS1, whereas TEAD1/2 loss expands PS2. This insight is not obtained from immunohistological analysis, likely due to the heterogeneity of the OLIG2^+^ASCL1^+^ state. That YAP/TAZ loss reduces PS1 is consistent with YAP/TAZ being predominantly expressed in APs and suggests that YAP/TAZ promote AP self-renewal and/or proliferation. Whether YAP/TAZ require TEAD for this function is unclear, as the effect of TEAD1/2 loss in PS2 might have masked an effect in PS1 by, for example, causing PS2 cells to revert back to PS1 cells as in the case of *Scalloped* mutants (see below); compensation by *Tead3* or *Tead4* is unlikely because they were not upregulated in *Tead1;2* dKO PS1 cells (data not shown).

TEAD loss and YAP/TAZ loss likely affect not only the number of cells belonging to a progenitor state but also the cell state (of the general sense) itself. In our “cluster tree” analysis, *Tead1;2* dKO progenitor cells switched between clusters upon a small change in clustering resolution more readily than WT progenitor cells did (data not shown), suggesting that the cell states of *Tead1;2* dKO cells are less well defined than that of WT cells. Furthermore, differential expression and GSEA analyses comparing dKO and control cells of the same progenitor state revealed many significantly changed gene sets (Supplemental Table S2 and data not shown), indicative of different biological activities—which constitute a cell state—between dKO and control cells in the same progenitor state. This type of analyses led us to discover that TEAD promotes subpallial progenitor lineage progression at least in part by inhibiting Notch signaling.

Based on the co-expression and physical interaction between TEAD and INSM1 in the subpallium and the phenotypic similarity between their mutants—both showed expansion of the OLIG2^+^ASCL1^+^ population, we suggest that TEAD acts with INSM1 to promote subpallial progenitor lineage progression. The physical interaction between orthologs of TEAD and INSM1 is evolutionarily conserved (Feng et al. 2013; Vissers et al. 2018; Guo et al. 2019). Our study demonstrates that the function of this transcription factor complex in regulating neurogenesis is also conserved. In *C. elegans*, this complex (EGL-44 and EGL-46) instructs cell-cycle exit during the lineage progression of a specific neuroblast (Feng et al. 2013) and controls cell fate in specific neurons (Wu et al. 2001). In *Drosophila*, this complex (Scalloped and Nerfin-1) maintains specific neuronal fate; in its absence, terminally differentiated medulla neurons in the optic lobes revert back to a neuroblast-like state (Vissers et al. 2018). Interestingly, Scalloped and Nerfin-1 maintains medullar neuron fate at least in part by repressing Notch signaling, which echoes our finding that TEAD1/2 promote subpallial progenitor lineage progression also by, at least in part, inhibiting Notch signaling. In both *C. elegans* and *Drosophila*, orthologs of TEAD and INSM1 only act together in specific contexts, while both playing additional roles independent of the other. During mammalian neural development, INSM1 has been shown to promote the generation and expansion of BPs in the cortex (Farkas et al. 2008), specify monoaminergic neuronal phenotypes in the hindbrain (Jacob et al. 2009), control the development of pituitary endocrine cells (Welcker et al. 2013), and consolidate outer hair cell fate in the cochlea (Wiwatpanit et al. 2018). It remains to be determined whether INSM1 functions with TEAD in any of these contexts.

In summary, we propose that during subpallial development, TEAD interacts with different partners in different progenitor subtypes/states to execute distinct functions (Fig. 7E). In APs, TEAD interacts with YAP/TAZ to maintain neuroepithelial structural integrity; whether TEAD also promotes self-renewal and proliferation with YAP/TAZ remains to be determined. In IPs, including SAPs and possibly a subset of BPs, TEAD interacts with INSM1 to promote lineage progression at least in part by inhibiting Notch signaling.

## Materials and Methods

### Mice

The *Nestin-Cre* line (Tronche et al. 1999) (#003771) and *Notch1^F^*allele (Yang et al. 2004) (#007181) was purchased from the Jackson Laboratory (JAX). *Tead2^F/F^* mice (Kaneko et al. 2007), *Yap^F/F^;Taz^F/F^*mice (Xin et al. 2011), and *Insm1^F/F^* mice (Wiwatpanit et al. 2018) were kindly provided by Drs. Melvin DePamphilis, Eric Olson, and Jaime García-Añoveros, respectively. To generate the *Tead1^F^* allele, *Tead1^tm1a^* (#050015-UCD) was obtained from Mutant Mouse Resource & Research Centers UC Davis and crossed with the *Rosa26-FLP* line (JAX #016226). Both *Tead1* and *Tead2* are on chromosome 7. A recombined chromosome 7 harboring both *Tead1^F^* and *Tead2^F^*alleles was generated by crossing *Tead1^F/+^;Tead2^F/+^*transheterozygous mice with wild-type mice and screening for progenies carrying both *Tead1^F^* and *Tead2^F^* alleles. Two *Vgll4^F^* lines were generated by using CRISPR-Cas9–mediated gene editing to insert two *loxP* sites flanking exon 2 in one line and exon 3 in the other. The corresponding deleted alleles (*Vgll4^Δ^*) were generated by crossing *Vgll4^F/F^* mice with *Sox2-Cre* (JAX #008454) females, which express *Cre* in their germline. All mice were maintained in a mixed genetic background. All animal procedures were approved by the Institutional Animal Care and Use Committee of St. Jude Children’s Research Hospital (SJCRH).

### Histology

Mouse brains were dissected in cold PBS, fixed overnight in 4% paraformaldehyde at 4°C, and washed in PBS. For paraffin sections, fixed brains were dehydrated in 70% ethanol overnight, embedded in paraffin, and sectioned at 5-μm thickness. For histological analysis, sections were rehydrated and stained with hematoxylin and eosin (H&E) or Luxol blue and Nissl. For frozen sections, fixed brains were cryoprotected in 30% sucrose overnight at 4°C, embedded in Tissue Freezing Medium (Thermo Scientific), and sectioned at 12–20-μm thickness. For immunostaining, sections were washed twice in PBS, blocked and permeabilized in PBS with 0.2% Triton X-100 (PBST) and 3% normal donkey serum for 1 h, and incubated with primary antibodies at 4°C overnight. Sections were then washed in PBST 3 times and incubated with appropriate fluorescence–conjugated secondary antibodies (Jackson ImmunoResearch Laboratories or Thermo Scientific Invitrogen) at 1:1000 dilution and DAPI for 2–3 h at room temperature. The following primary antibodies were used: Alexa Fluor 647-Sox2 (BD Biosciences, #562139, 1:100), BrdU (Abcam ab6326, 1:100), β-catenin (BD Biosciences #610153, 1:500), TBR2/EOMES (eBioscience, #14-4875, 1:100), phospho-Histone H3 (Ser10) (Cell Signaling Technologies (CST) #9706, 1:500), INSM1 (Santa Cruz Biotechnologies sc-271408 AF488, 1:50), Ki67 (Invitrogen #14-5698-82, 1:1000), MAFB (Sigma HPA005653, 1:500), ASCL1 (Abcam ab211327, 1:500), PKCζ/aPKC (Santa Cruz sc-216, 1:1000), SOX10 (Santa Cruz sc-17342, 1:250), OLIG2 (R&D, af2418, 1:250), TEAD1 (CST #12292, 1:250), TUJ1 (BioLegend #801202, 1:100), YAP/TAZ (CST #8418, 1:500), and ZO-1 (Invitrogen #339188, 1:500).

### Image Quantifications

All images used for cell counting were acquired using a Zeiss LSM 780 confocal or Zeiss Apotome microscope with a 40x objective lens. For each genotype, 3 animals with 1–3 sections of comparable positions per animal were quantified. Areas of interest were outlined and isolated in ImageJ. Nucleus segmentation was carried out using the machine-learning–based segmentation method StarDist (https://github.com/stardist/stardist), which was first trained using manually annotated datasets. Fluorescent label intensity of each channel over nuclear ROIs was calculated by a custom multi-channel intensity quantification algorithm created in OMERO (https://www.openmicroscopy.org/omero/scientists/). Thresholds for signal positivity were manually determined by inspecting each image. Mitosis location, MAFB-labeled cells and OLCs were counted manually in OMERO. For TUJ1 analysis, areas of interest were outlined and the percentage of TUJ1^+^ regions were measured using the Olympus CellSens Dimension analysis program (Olympus). Image area and ventricular length were measured with Zeiss Zen 3.9 imaging software. Data points in all quantification graphs correspond to individual animals, which are the average of 1–3 sections per animal.

### Western blot

Mouse brains were dissected and total cell lysates were prepared in 20 mM HEPES (pH 7.4), 150 mM NaCl, 2% SDS and 5% glycerol supplemented with AEBSF and Halt protease and phosphatase inhibitors (Thermo). Protein concentrations were measured by BCA assay (Thermo). Twenty μg of protein per lane were subjected to SDS-PAGE and probed with primary antibodies and HRP-conjugated secondary antibodies. The following primary antibodies were used: β-actin (Ambion #AM4302, 1:40,000), GAPDH (CST #2118S, 1:3000), FLAG (CST #2368S, 1:1000), INSM1 (Santa Cruz sc-271408, 1:500), TEAD1 (CST #12292, 1:500), pan-TEAD (CST #13295, 1:500), and VGLL4 (Thermo #PA5-58303, 1:50). Each lane corresponds to an individual embryo.

### Co-immunoprecipitation

Plasmids encoding FLAG-tagged proteins were transfected into HEK293T cells plated on 10-cm plates using Lipofectamin 2000. Co-immunoprecipitation was performed 48 h later using Nuclear Complex Co-IP kit (Active Motif #54001) according to kit instruction, using IP Low buffer supplemented with additional detergent to 1% final concentration and 10 µl of Pan-TEAD (CST #13295, 33 µg/ml) or 5 µl of FLAG-M2 (Sigma #F1804, 1 µg/µl) antibody.

### Proximity ligation assay

The assay was performed according to Duolink kit (Millipore Sigma DUO92103) protocol. The following primary antibodies were used: TEAD1(CST #12292, 1:250), YAP (Sigma WH0010413M1, 1:1000), INSM1 (Santa Cruz Biotechnologies sc-271408 X, 1:50), Ki67 (BD Pharmigen #558615, 1:250) and Ki67 (Vector VP-RM0, 1:250). Images used for quantification were acquired using a Zeiss LSM 780 confocal microscope and a 40x objective lens. The number of PLA spots per image were quantified using OMERO. Each interaction was quantified in 3 embryos. At least 4 images from subpallial sections of comparable positions were quantified for each embryo.

### Single cell RNA sequencing

MGEs were obtained by first extracting the embryonic brain and detaching the telencephalon, followed by removing the dorsal telencephalon and finally dissecting the MGE from the rest of the subpallium in cold PBS. Both MGEs from the same embryo were dissociated using either Stem-Pro Accutase (Thermo Scientific A1110501) or a papain dissociation kit (Worthington LK003150) for less than 5 min. Cells were filtered through Flowmi cell strainers (SP Bel-Art #13680040). Cell viability was analyzed with a Countess cell counter and only samples with a viability of >90% were processed for scRNAseq. A 3’ scRNAseq kit from 10x Genomics was used according to kit instructions. Up to 10K cells were loaded per each bead mix.

### Single cell RNA sequencing data analysis

General quality control and filtering: Sequences from each individual Illumina sequencing dataset were demultiplexed using bcl2fastq v2.20.0.422 (Illumina). Sequencing reads were processed using 10X Genomics Cell Ranger version 7.0.0, with reads mapping to the mouse reference genome mm10 version 2020-A (10x Genomics). Quality control filtering, clustering, dimensionality reduction, visualization, and differential gene expression were performed using Seurat v4.0.4 with R v4.1.0 (Stuart et al. 2019). Each dataset was initially filtered so that genes that were expressed in at least three or more cells and cells that expressed at least 200 genes were retained when reading in the data. Then cells were excluded if they: (1) expressed greater than 10% of unique transcripts derived from mitochondrial genes, (2) unique transcripts derived from mitochondrial genes expressing more than 3 median absolute deviations (MADs) from the median number of unique transcripts derived from mitochondrial genes, (3) had fewer than 1,500 genes expressed, or (3) had UMI counts below 5,000. Afterward, cells with more than 3 MADs from the median number of genes expressed were removed.

Dataset integration processing: Datasets were individually log-normalized using the *NormalizeData* function with default parameters. We calculated 3,000 features that exhibit high cell-to-cell variation in the dataset using the *FindVariableFeature* function. Next, we scaled the data by linear regression against the number of reads using the *ScaleData* function with default parameters. The variable genes were projected onto a low-dimensional subspace using principal component analysis using the *RunPCA* function with default parameters. The number of principal components (Npcs) were selected based on inspection of the plot of variance explained (Npcs = 30). Datasets were integrated using Harmony integration with default parameters. A shared-nearest-neighbor graph was constructed based on the Euclidean distance in the low-dimensional subspace using the *FindNeighbors* function with dims = 1:30 and default parameters. Integrated datasets then underwent non-linear dimensional reduction and visualization using UMAP. Clusters were identified using a resolution of 0.3. Gene signature scores were calculated using the *AddModuleScore* function with our lists of progenitor genes and neuron genes (Supplemental Table S1). Cell cycle phase was inferred using the S- and G2M-phase gene lists from Tirosh, et al 2015 using the *CellCycleScoring* command in Seurat.

Progenitor cluster integration processing: Clusters with average progenitors scores above zero (WT dataset clusters 2,3,6, WT/KO dataset clusters 2,3,5) were subset out and re-analyzed following the above processing steps except that (1) cells with fewer than 2,500 genes expressed were removed and that (2) a list of 742 transcription factor genes (Supplemental Table S1) was used as the features to project onto a low-dimensional subspace using principal component analysis using the *RunPCA* function with default parameters. Gene signature scores were calculated using the *AddModuleScore* function with our lists of VZ genes and SVZ-MZ genes (Supplemental Table S1). Cell cycle phase was inferred as described above. Pseudotime analysis was conducted using Monocle3 (v1.3.1) with default parameters. Trajectory starting points were manually selected based on the maximum expression of VZ markers. Cluster tree was generated by stepwise (0.025) increasing clustering resolution using progenitor cells from the WT dataset (WT dataset clusters 2,3,6), which identified 0.25 as the optimal resolution. To assign progenitor state annotation to mutant cells, we used the *FindIntegrationAnchors* function in Seurat to detect integration anchors, using the first 30 PCA dimensions between the WT_progenitors_dataset (reference) and the All_progenitors_dataset (query). These anchors were then used to transfer progenitor state labels from the WT_progenitors_dataset to the All_progenitors_dataset. The transfer efficiency was validated by examining the clustering consistency of shared cells between the two datasets, achieving ∼93% agreement. The *propeller* function from the speckle_v_0.0.3 package (Phipson et al. 2022) was used to evaluate statistical differences in cell proportions between control and mutants. Pseudotime density plot was generated using the *geom_density* function in ggplot. Differential expression analysis was performed using the *getDE* function in scMINER (Ding et al. 2023) (https://jyyulab.github.io/scMINER). Gene set enrichment analysis (GSEA) was performed using the *funcEnrich.Fisher* function and visualized with the *draw.funcEnrich.cluster* function in NetBID2 (Dong et al. 2023) (https://jyyulab.github.io/NetBID). Selected gene sets were visualized with the *draw.GSEA* function in NetBID2.

### Calculation of in vivo cell cycle parameters

Pregnant dams were injected intraperitoneally with EdU (10 μg/g body weight) and followed by BrdU (50 μg/g body weight) 1.5 h later. Dams were euthanized 30-min following BrdU administration, and embryos were collected and processed for cryosectioning. Sections were blocked in PBST containing 3% normal donkey serum, incubated in primary antibodies against ASCL1 and OLIG2 at 4°C overnight, washed in PBST, and incubated in fluorescence-conjugated secondary antibodies for 2 h at room temperature. After 3 washes in PBST, sections were fixed in 4% paraformaldehyde for 10 min, followed by 3 washes in PBST. Sections were incubated in 1N HCl at 45°C for 30 min then washed in PBST. EdU was detected by the Click-it Plus EdU Alexa Fluor 488 Imaging Kit (Invitrogen #C10337). Sections were then washed in PBST, blocked, incubated in BrdU antibody (Abcam #ab6326, 1:100) overnight at 4°C and secondary antibody for 2 h at room temperature. Z-stack images (5) were acquired on a Zeiss LSM 780 confocal microscope, merged in ImageJ, and quantified with OMERO.

Cell cycle parameters were calculated based on the method described (Quinn et al. 2007). The following populations were quantified using OMERO: Cycling fraction (Cycling_fraction_): total number of cells in each progenitor subtype defined as OLIG2^+^ASCL1^−^, OLIG2^+^ASCL1^+^, and OLIG2^−^ASCL1^+^; S-phase fraction (S_fraction_): the number of progenitor cells in the S phase, defined as EdU^+^BrdU^+^ cells; Leaving fraction (L_fraction_): the number of progenitor cells that have left the S phase, defined as EdU^+^BrdU^−^ cells. S-phase length (T_S_) and total cell-cycle length (T_C_) were then calculated using the following equations:

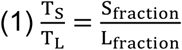

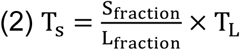

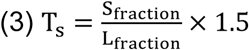

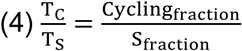

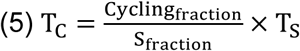

### FlashTag labeling of neural progenitor cells in vivo

The procedure was performed according to the protocol described (Govindan et al. 2018). Pregnant mice carrying E12.5 embryos were deeply anesthetized with 2.5–3% of isoflurane and the uterine horns exposed. Approximately 300 nL of 10 mM carboxyfluorescein succinimidyl ester (CFSE, Invitrogen C34554) in DMSO was injected into a lateral ventricle of the embryo brain using a pulled glass micropipette. Embryos were returned to dam and allowed to resume normal development before being harvested for immunostaining.

### RNAscope in situ hybridization

Tissue sections were processed according to the kit instructions for frozen samples (Advanced Cell Diagnostics (ACD) RNAscope Multiplex FL v2 #323100). Tissue was permeabilized with Protease III for 20 min at 40°C. Sections were incubated with a probe mixture targeting MmHes5 (ACD #400991, C2) and MmNotch1 (ACD #404641, C4), each at a 1:100 dilution.

Fluorescent images were acquired using a Zeiss Apotome microscope. Laser power and exposure time was adjusted to ensure below saturation exposure. Quantification was performed in ImageJ. The mean fluorescence intensity (total fluorescence signal divided by area) of *Hes5* and *Notch1* was measured in the medial ganglionic eminence (MGE) and a cortical VZ region within the same section. Because the cortex of *Tead1;2* dKO mice did not shown obvious morphological phenotype, the cortex was used as an internal control. The average mean intensity values from the left and right MGE were normalized to the mean intensity values in the cortex.

### Statistical analysis

Statistical analyses of immunostaining and RNAscope experiments were done using GraphPad Prism software. Statistical significance was assessed by unpaired two-tailed t-test. *P* < 0.05 was considered a significant difference. Individual animals are defined as a biological replicated. All values represent individual animals mean ± SEM (standard error of the mean). The statistical test used and the P values are indicated in figures or figure legends. Statistical analyses of scRNA-seq data were performed using designated functions in *propeller* (cell proportions), scMINER (differential gene expression), and NetBID2 (gene set enrichment analysis).

## Acknowledgements

We thank members of the Cao lab for suggestions and technical help; Aaron Taylor and George Campbell for help in image acquisition; Abbas Shirinifard for help in image quantification; Yiping Fan and Hongjian Jin for help in scRNA-seq data analysis; Shondra Miller, Valerie Stewart, and members of the Center for Advanced Genomic Editing and of the Neuroembryology core at St. Jude Children’s Research Hospital (SJCRH) for generating the *Vgll4^F^* alleles; Melvin DePamphilis, Eric Olson, and Jaime García-Añoveros for sharing mouse lines. This work was supported by National Institute of Health grant R01NS119760 (to X.C. and J.Y.), P30CA021765 (to SJCRH Comprehensive Cancer Center core facilities), and American Lebanese Syrian Associated Charities (ALSAC).

## Author Contributions

X.C. conceived and supervised the project; C.P and A.L. performed the majority of the experiments and data analysis; A.L., V.T., C.R., A.M., J.Y. and J.Y performed computational analysis of scRNA-seq data; J.P. helped with mouse management and Western blotting; R.D. helped with histological analysis. X.C. wrote the paper with input from other authors.

## Supplemental Material

Supplemental Figures S1 and S2. Additional data related to Figure 1.

Supplemental Figure S3. Additional data related to Figure 4.

Supplemental Figures S4 and S5. Additional data related to Figure 7.

Tables S1. Gene lists used for scRNA-seq data analysis. Related to Figure 3.

Tables S2. Differential expression and GSEA analyses of PS2 cells. Related to Figure 6.

---

**Supplemental Figure S1.**
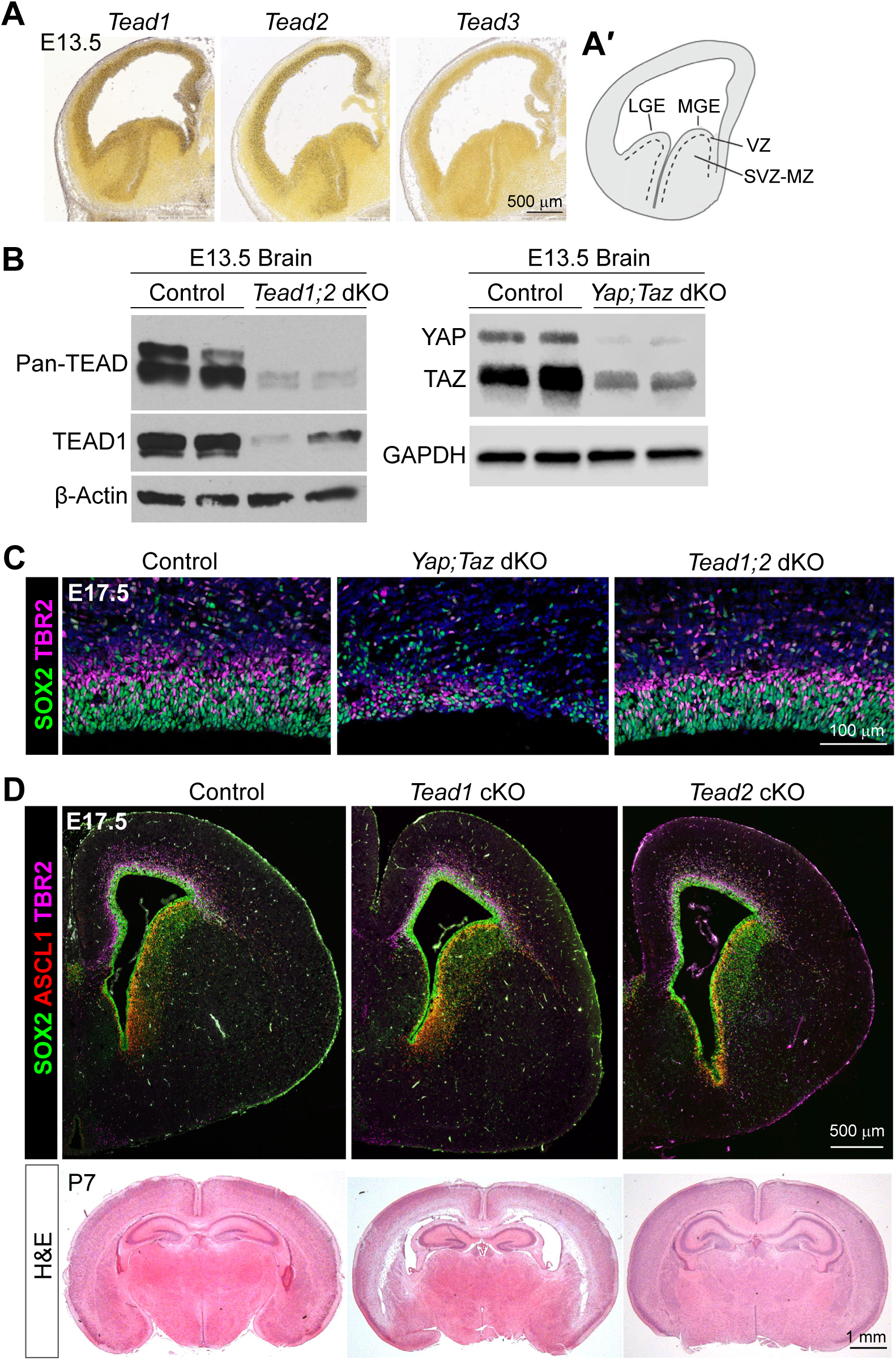
Expression patterns of *Tead* genes in the developing telencephalon and the gross phenotypes of *Yap;Taz* and *Tead* knockout mice. (A) RNA in situ hybridization images of *Tead1*–*3* genes (Allen Developing Mouse Brain Atlas). (A′) A schematic of the E13.5 telencephalon showing subpallial regional organization. LGE, lateral ganglionic eminence; MGE, medial ganglionic eminence; VZ, ventricular zone; SVZ, subventricular zone; MZ, mantle zone. (B) Western blot analysis of total brain lysates from no-*Cre* control and *Nestin-Cre*–mediated *Tead1* and *Tead2* double knockout (*Tead1;2* dKO) and *Yap* and *Taz* (*Yap;Taz*) dKO mice. Note that a TEAD2 specific antibody was not available to us. Nevertheless, a pan-TEAD antibody showed that the levels of TEAD proteins were strongly reduced in *Tead1;2* dKO brains. (C) Immunostaining showing the neocortex of control and dKO mice, focusing on the VZ and SVZ regions. Blue color is DAPI signal. (D) Immunostaining and H&E staining showing the gross brain morphology of *Nestin-Cre*–mediated *Tead1* and *Tead2* single conditional knockout (cKO) mice at E17.5 and postnatal day (P) 7.

**Supplemental Figure S2.**
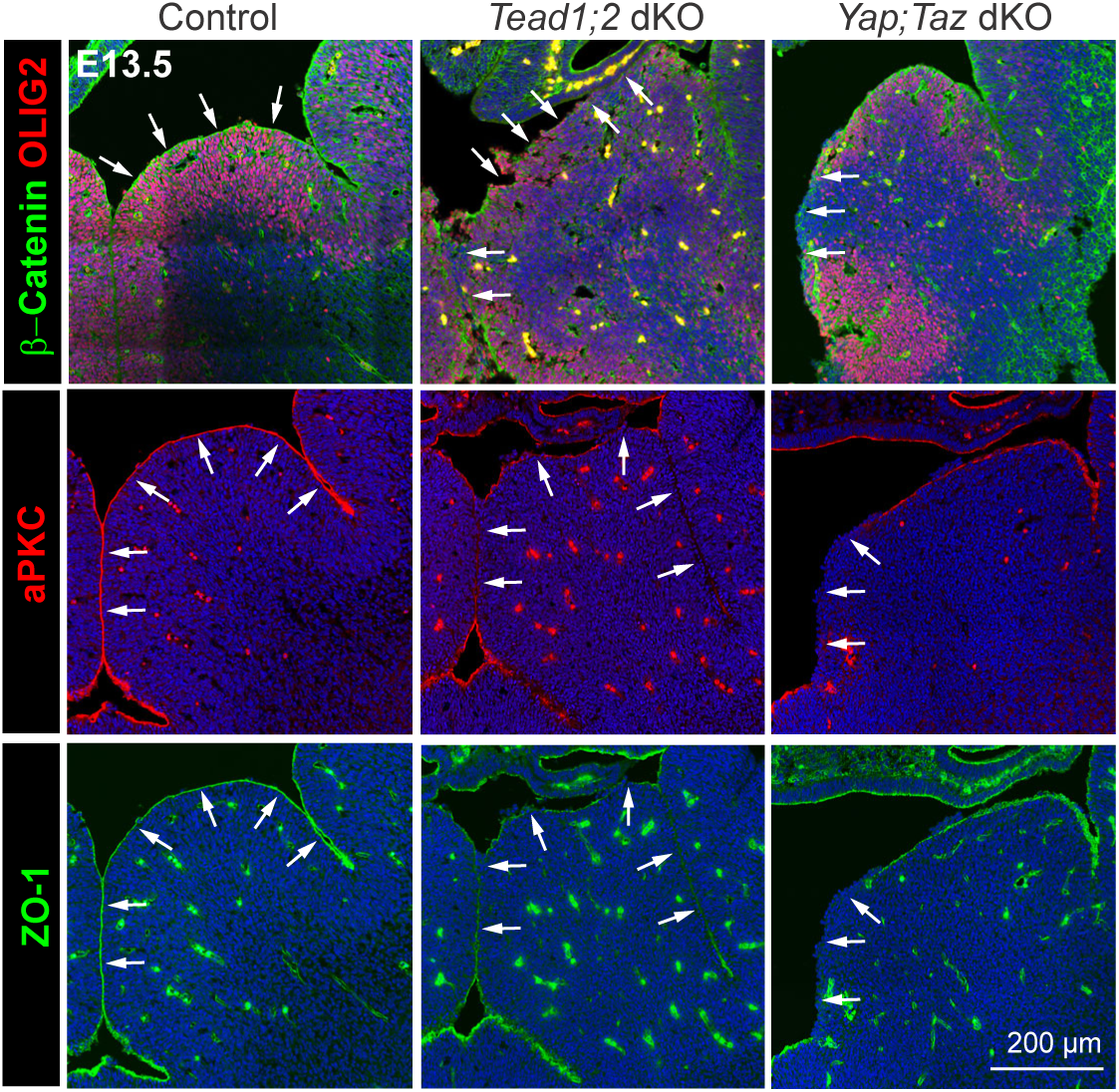
TEAD1/2 loss and YAP/TAZ loss similarly disrupt subpallial neu-roepithelial apical juctions. Immunostaining with apical junction markers β-catenin, aPKC, and ZO-1 showing the medial ganglionic eminence. Arrows: apical junctions. Blue color is DAPI signal.

**Supplemental Figure S3.**
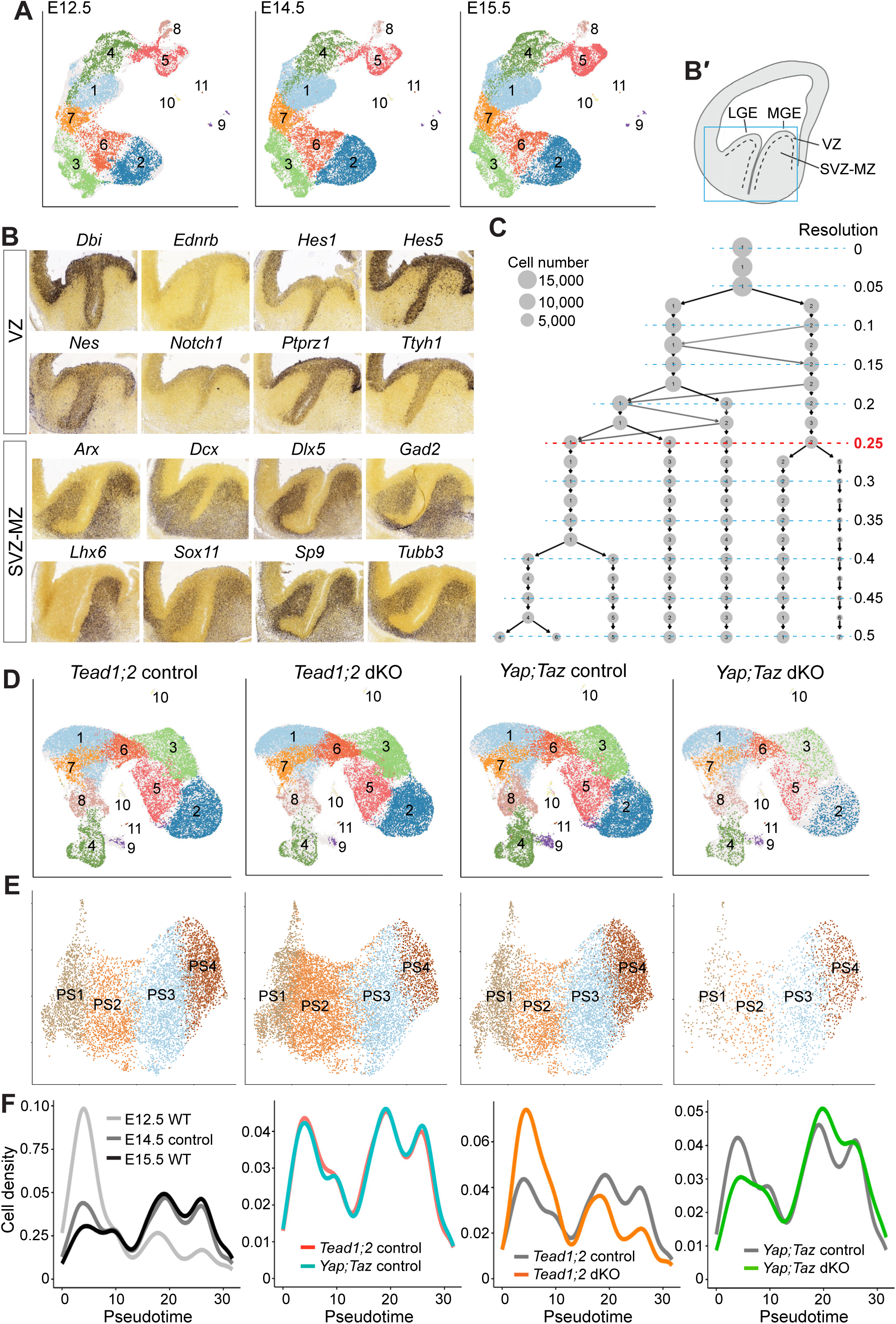
Single-cell RNA sequencing analysis reveals distinct impacts of TEAD1/2 loss and YAP/TAZ loss on MGE neural progenitor lineage progression. (A) Clustering of WT MGE cells based on highly variable genes (HVG) visualized by UMAP. E12.5 WT, 11,929 cells after quality control; E14.5 *Tead1;2* control, 15,124 cells; E15.5 WT, 15,823 cells. (B) Expression patterns (from Allen Developing Mouse Brain Atlas) of selected genes in the VZ and SVZ-MZ gene lists. (B′) A schematic of the E13.5 telencephalon showing subpallial regional organization. VZ, ventricular zone; SVZ, subventricular zone; MZ, mantle zone. (C) Cluster tree analysis of WT progenitor cells showing changes in cluster composition with stepwise increase of clustering resolution. (D) Clustering of E14.5 control and dKO MGE cells based on HVG visualized by UMAP. *Tead1;2* control, 15,124 cells; *Tead1;2* dKO, 18,406 cells; *Yap;Taz* control, 18,041 cells; *Yap;Taz* dKO, 6,361 cells. (E) Progenitor state annotation of progenitor cells visualized by UMAP using the list of transcription factors for feature selection. *Tead1;2* control, 7,172 cells; *Tead1;2* dKO, 9,648 cells; *Yap;Taz* control, 8,299 cells; *Yap;Taz* dKO, 1,954 cells. (F) Pseudotime density plot of progenitor cells.

**Supplemental Figure S4.**
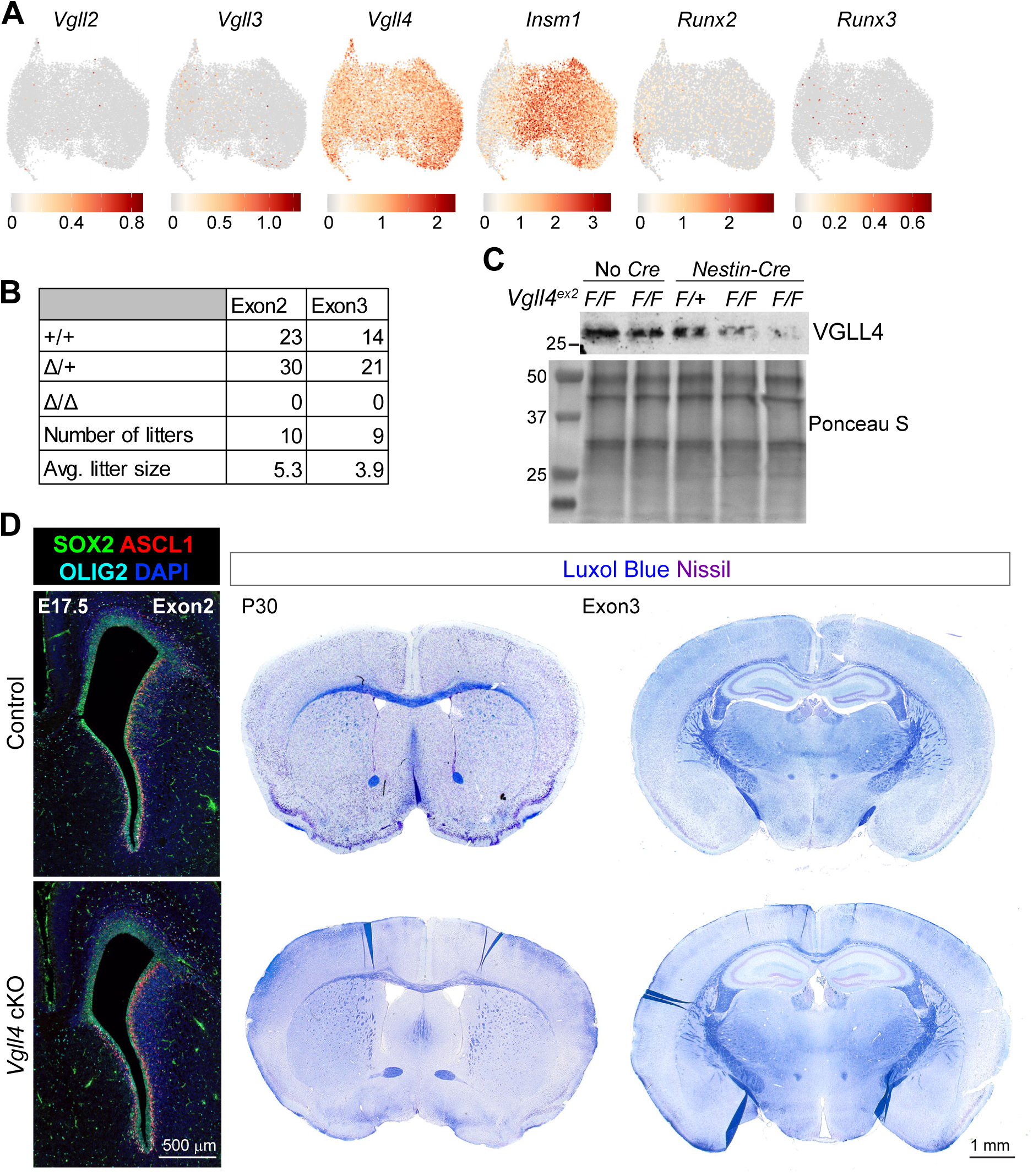
Loss of VGLL4 does not notably perturb brain development. (A) ScRNA-seq expression levels of genes encoding proteins known to interact with TEAD in WT progenitor cells visualized by UMAP. (B) Number of live animals genotyped from *Vgll4*^Δ/+^ x *Vgll4*^Δ/+^ breedings. Tissue biopsies were collected between P3 to P7. (C) Western blot of total brain lysates from E14.5 no-*Cre* control and *Nestin-cre*-mediated *Vgll4* conditional KO (cKO) mice deleting exon 2. (D) Gross morphology of control and *Vgll4* cKO brains. E17.5 images are obtained from exon2 cKO line. P30 images are from exon3 cKO line.

**Supplemental Figure S5.**
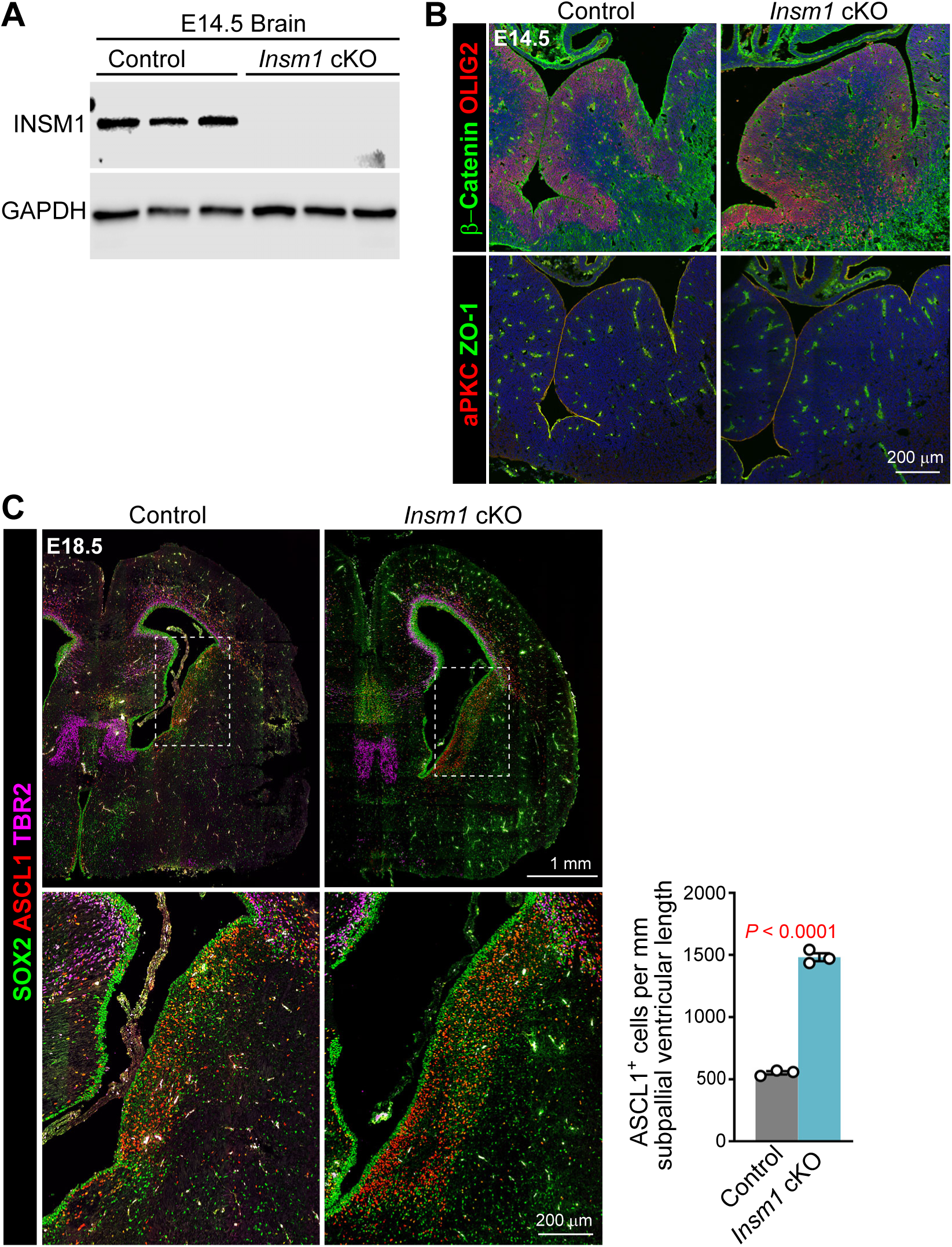
Characterization of *Insm1* cKO mice. (A) Western blot of total brain lysates from no-*Cre* control and *Nestin-cre*-mediated *Insm1* conditional KO (cKO) mice. (B) Immunostaining for neural progenitor (OLIG2) and apical junction (β-catenin, aPKC, ZO-1) markers, showing MGE regions. Blue color is DAPI signal. (C) Immunostaining of neural progenitor cells and quantification of ASCL1-labeled cells in subpallial VZ-SVZ regions. Areas in dashed boxes are enlarged in the images below. Values are mean ± SEM. Two-sided unpaired *t*-test. Each data point represents an individual animal.

